# CMV-specific clonal expansion of Th1, GZMK⁺ CD8⁺, and TEMRA T cells revealed by human PBMC single cell profiling

**DOI:** 10.1101/2025.06.24.661167

**Authors:** Titas Grabauskas, Luke Trinity, Chris P. Verschoor, Djamel Nehar-Belaid, Radu Marches, Giray Eryilmaz, Avinash S. Mahajan, Sathya Baarathi, Asa Thibodeau, Emilie Picard, Chia-Ling Kuo, Kenneth E. Schmader, Cathleen Colon-Emeric, Heather E. Whitson, Silke Paust, Adolfo García-Sastre, Gur Yaari, Jacques Banchereau, George A. Kuchel, Duygu Ucar

## Abstract

Cytomegalovirus (CMV) is a common herpesvirus that establishes lifelong latency and becomes increasingly prevalent with age. We systematically characterized CMV-associated immune remodeling by analyzing six human cohorts (two newly built) using single-cell RNA sequencing, T cell receptor (TCR) sequencing, and flow cytometry. Beyond the well-known expansion of CD4⁺/CD8⁺ TEMRA, adaptive NK, and γδ T cells, CMV(+) adults exhibited increased frequencies of GZMK⁺ CD8⁺ T cells and atypical B cells, alongside a reduction of CD56^dim^ NK cells. Longitudinal profiling of an individual who seroconverted revealed rapid CMV-driven shifts in circulating immune cell frequencies. Single-cell TCR data analyzed using a large database of CMV-associated clones combined with predictive modelling (CMVerify), identified novel CMV-specific clonal expansions reproduced across two independent cohorts. In the CD8⁺ lineage, CMV-specific clones were enriched in GZMK⁺ CD8⁺ and CD8⁺ TEMRA cells, while in the CD4⁺ lineage, Th1 cells showed clonal expansion alongside CD4⁺ TEMRA cells. This integrative study revealed how latent CMV alters the cellular and clonal landscape, defining GZMK⁺ CD8⁺ and Th1 cells as newly recognized elements of response to CMV in humans.

## Introduction

The immune system undergoes significant changes with aging, characterized by a decline in naïve cell populations, expansion of memory and terminally differentiated T and B cells, and increased pro-inflammatory activity^1,2^. These changes contribute to weakened immune responses and increased susceptibility to disease^1,3,4^. However, the trajectory of immune aging varies widely between individuals^5^, leading to differences in immune health, vaccine efficacy, and therapeutic responses^6–8^. Twin studies have shown that immune variability increases with age and non-heritable environmental factors account for a substantial proportion of this inter-individual variation^9^. Notably, cytomegalovirus (CMV) infection has emerged as a key environmental factor, as monozygotic twins discordant for CMV exhibited significant divergence in immune cell traits^9,10^.

CMV is a DNA herpesvirus transmitted through bodily fluids^11^, that establishes lifelong latency in hematopoietic stem and myeloid progenitor cells^12^. It is one of the most prevalent chronic infections worldwide, with seropositivity increasing with age^13^. According to the CDC, over 50% of adults in the U.S. are infected with CMV by age of 40^14^. Although typically asymptomatic in healthy individuals, latent CMV has been associated with more severe outcomes of SARS-CoV-2^15^ and tuberculosis infections^16^. Conversely, CMV seropositivity has also been linked to improved survival following anti-PD-1 checkpoint blockade in older melanoma patients^17^, and enhanced influenza vaccine responses in healthy young adults^18^. Despite its widespread prevalence and known impact on immune heterogeneity, CMV serostatus is often unreported in single-cell studies of immune aging^2,19,20^, limiting our ability to distinguish age-related changes from those associated with CMV.

A hallmark of CMV seropositivity is the expansion of effector memory and terminally differentiated (TEMRA) CD4⁺ and CD8⁺ T cells, observed in both young and older adults^21–25^. This phenomenon, known as “memory inflation,” reflects the long-term accumulation of CMV-specific clones, particularly within the TEMRA subset (CD45RA⁺CCR7⁻)^26,27^, which can comprise up to 10–30% of the total memory T cell pool^26,28^. Clonal expansion of CMV-specific CD8^+^ T cels reduces the overall T cell receptor (TCR) repertoire diversity^29–31^. While TEMRA cells remain functional—typically expressing *GZMB*, *PRF1*, and *KLRG1*^2,32^ and exhibiting potent cytotoxic capacity^22,26,28^—they have limited proliferative potential^33^.

Beyond conventional αβ T cells, Vδ1 γδ T lymphocytes also expand in CMV-positive individuals and display adaptive-like features, including clonal expansion and cytotoxic responses to infected cells^34^. Latent CMV infection has been associated with expansion of adaptive NK cells expressing NKG2C^35,36^, an activating receptor that recognizes CMV-derived peptides and restricts viral spread^37,38^. These NKG2C⁺ NK cells persist long-term and display features of immunological memory^38,39^.

While these critical insights were derived largely within specific lineages using protein-level data, the recent availability of high-resolution single-cell multi-omic datasets (e.g., RNA-seq, TCR-seq) across independent cohorts and age groups has now enabled a systematic, unbiased dissection of CMV-associated immune remodeling at unprecedented depth. For this purpose, we took advantage of publicly available single-cell datasets and built two new cohorts of our own (see Methods), uncovering novel cellular and clonal associations with CMV, including clonal expansions in GZMK⁺ CD8⁺ and Th1 cells.

### Study design: newly built older adult cohort for CMV transcriptional signature discovery

We recruited two cohorts of healthy older adults during 2017-2018 from communities surrounding Hartford, Connecticut, USA, and Sudbury, Ontario, Canada. Cohort 1 comprised 36 healthy older adults (n=17 CMV-positive hereinafter referred to as CMV(+), n=19 CMV-negative referred to as CMV(-)), whose peripheral blood mononuclear cells (PBMCs) were profiled using single-cell RNA sequencing (scRNAseq) and flow cytometry (Fig. 1a). Cohort 2 included 63 older adults (n=40 CMV(+), n=23 CMV(-)) whose PBMCs were profiled using flow cytometry alone^40,41^ (Fig. 1a). Participants were free from confounding autoimmune diseases, were not undergoing treatments that might influence immune function and composition (e.g., immunotherapy) and were not recently vaccinated before blood collection. CMV serostatus was determined in serum using ELISA for CMV IgG antibodies.

**Fig. 1.**
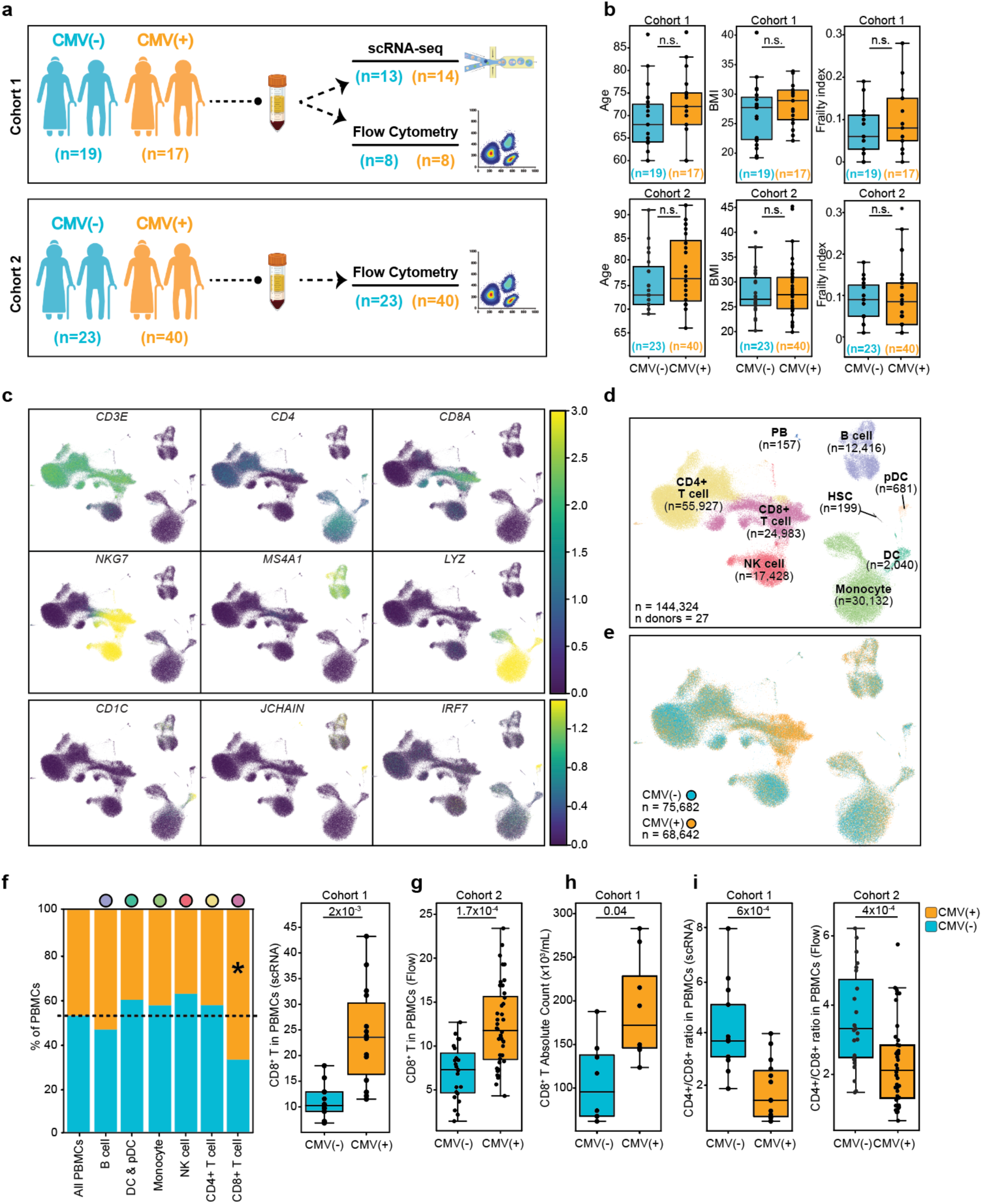
Study design and CD8^+^ T cell subset expansion associated with CMV. **a,** Schematic representation of the study design with two cohorts. *Top panel*: Discovery Cohort included n=19 CMV(-) and n=17 CMV(+) older adults. *Lower panel*: Validation Cohort included n=23 CMV(-) and n=40 CMV(+) older adults. **b,** Distribution of age, BMI, and frailty index for CMV(-) and CMV(+) groups in Discovery Cohort (*top panel*) and Validation Cohort (*bottom panel*). n.s. indicates not significant. **c,** Uniform manifold approximation and projection (UMAP) representing PBMCs from Discovery Cohort showing expression of subset-specific marker genes. **d,** 9 subsets (n=144,324) colored by immune cell annotations. NK, natural killer; pDC, plasmacytoid dendritic cell; PB, plasmablast; HSC, hematopoietic stem-cell. **e,** UMAP colored for CMV serostatus. **f,** *Left*: Immune cell frequencies in CMV(-) and CMV(+). Cell types with significant frequency differences between CMV(-) and CMV(+) groups are marked with an asterisk (*; P-adjusted [Benjamini-Hochberg] < 0.05). *Right*: Individual-level quantification of CD8^+^ T cells within PBMCs in CMV(-) (n=13) and CMV(+) (n=14). **g,** CD8^+^ T cell frequency within PBMCs quantified by flow cytometry in Validation Cohort: CMV(-) (n=23) and CMV(+) (n=40). **h,** Absolute count of CD8^+^ T cells quantified by flow cytometry in Discovery Cohort: CMV(-) (n=8) and CMV(+) (n=8). **i,** *Left*: CD4^+^ to CD8^+^ T cell frequency ratios quantified *via* scRNA-seq in Discovery Cohort: CMV(-) (n=13) and CMV(+) (n=14). *Right:* CD4^+^ to CD8^+^ T cell frequency ratios quantified by flow cytometry in Validation Cohort: CMV(-) (n=23) and CMV(+) (n=40). Box plots display the median and IQR (25–75%), with whiskers representing the upper and lower quartiles ±1.5× IQR. Each dot represents an individual sample. Mann-Whitney-U test (two-sided) was used to compare cell counts between CMV(-) and CMV(+) (**b, f, g, h, i**). All *P*-values are shown after adjusting for multiple hypothesis testing using Benjamini-Hochberg at false discovery rate (FDR) < 0.05.

The median age of Cohort 1 was 71 years old, including men (n=17) and women (n=19). CMV(+) and CMV(-) groups were comparable in age (p=0.90), body mass index (BMI; median 27.8, p=0.51), and frailty index^42^ (median 0.085, p=0.50) (Fig. 1b, Extended Data Fig. 1a, and Supplementary Table 1). Other characteristics, including race, cancer history, and medication use, were also evenly distributed between the two groups (Supplementary Table 1).

Cohort 2 had a median age of 75 years, including men (n=24) and women (n=39). Similar to Cohort 1, the CMV(+) and CMV(-) groups were comparable in age (p=0.11), BMI (median 27.1, p=0.93), and frailty index^42^ (median 0.09, p=0.88) (Fig. 1b and Supplementary Table 1). Race and cancer history were also evenly distributed between the groups (Supplementary Table 1).

### Frequencies of memory CD8^+^ T and γδ T cells are higher in CMV(+) older adults

PBMCs from Cohort 1 were profiled using scRNAseq, capturing 144,324 cells (n=68,642 CMV(+), n=75,682 CMV(-)) after quality control (Fig. 1c,d, and Extended Data Fig. 1b). As a group, CMV(+) individuals had higher frequency of CD8⁺ T cells compared to CMV(-) individuals (25% vs 11% of PBMCs; p=0.0012; Fig. 1e,f and Extended Data Fig. 1c), confirmed by flow cytometry in Cohort 2 (p<0.001; Fig. 1g and Extended Data Fig. 1d). Absolute CD8⁺ T cell counts were 1.78-fold higher in CMV(+) individuals (p=0.04; Fig. 1h), and the CD4⁺/CD8⁺ T cell ratio was lower in CMV(+) older adults in both cohorts (Discovery: p=0.006; Validation: p<0.001) (Fig. 1i and Extended Data Fig. 1d).

Clustering of CD8^+^ T cells (n=24,983) (Fig. 2a and Extended Data Fig. 2a) identified seven subsets: (i) naïve (*CCR7^+^*, *SELL^+^*, *LEF1+*), (ii) central memory (CM; *CCR7*^+^), (iii) mucosal-associated invariant T (MAIT; *ZBTB16^+^*, *KLRB1^+^*), (iv) GZMK^+^ CD8^+^ (*CCR7*^-^, *GZMK*^+^), (v) TEMRA (*GZMB^+^*, *KLRD1^+^)*, (vi) *GZMK*^+^ *γ*δ (*TRDC^+^, GZMK^+^*), and (vii) *GZMB*^+^ *γ*δ (*TRDC^+^, GZMB^+^*) cells. All subsets, except MAIT cells, were significantly altered with CMV positivity. GZMK⁺ CD8⁺ (p<0.001), TEMRA (p<0.001), and GZMB⁺ γδ (p<0.001) were subsets with the highest increase in frequency in CMV(+) older adults (Fig. 2b,c and Extended Data Fig. 2b).

**Fig. 2.**
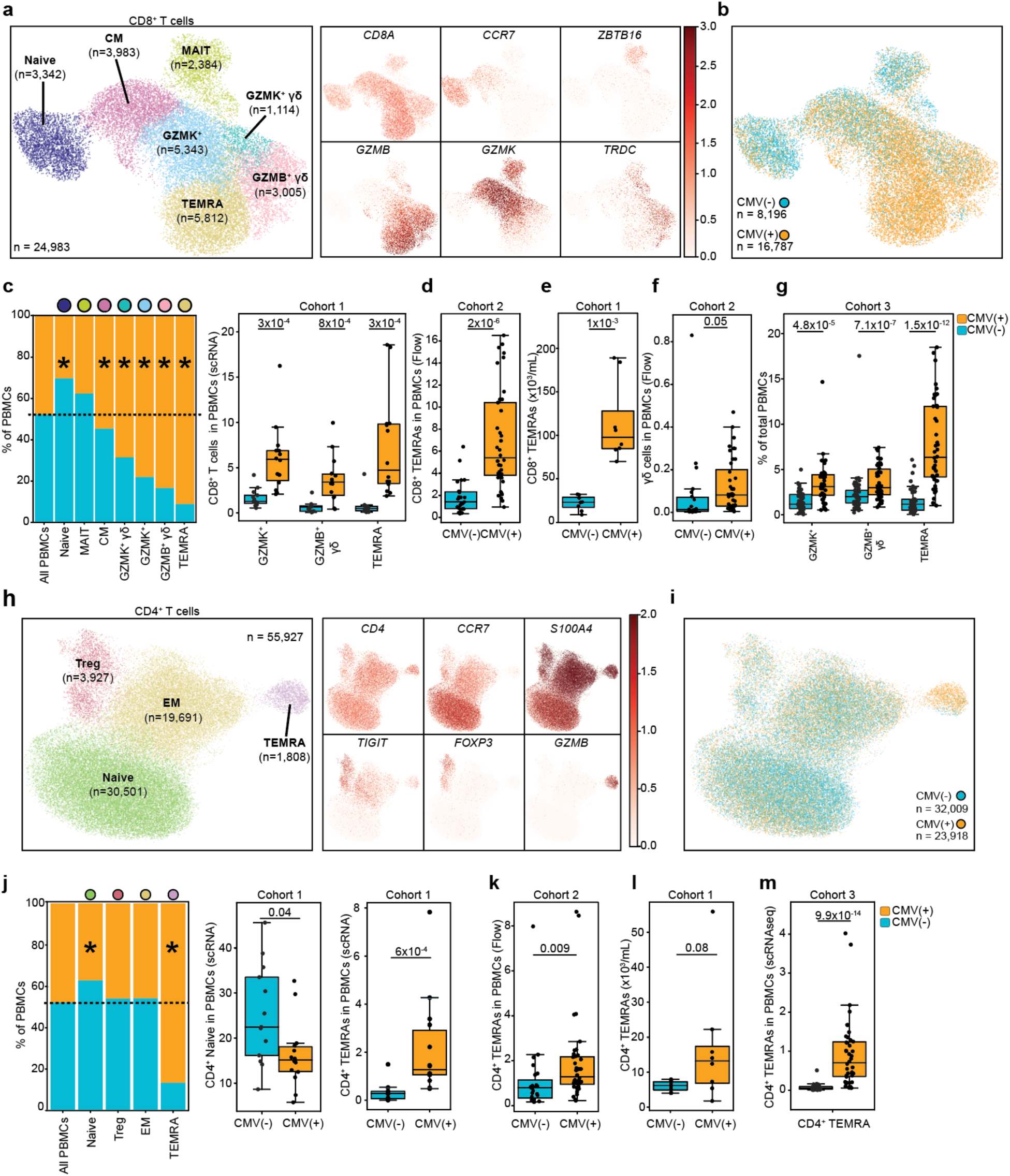
Expansion of memory CD8^+^ T, γδ T, and CD4^+^ TEMRA cell subsets associated with CMV positivity. **a,** *Left:* UMAP representing 7 subsets from CD8^+^ T cells (n= 24,983). CM, central memory; MAIT, mucosal associated invariant T; TEMRA, terminally differentiated effector memory. *Right:* UMAPs of CD8^+^ T cells showing expression of select marker genes. **b,** UMAP color coded for CMV serostatus. **c,** *Left:* CD8^+^ T cell subset frequencies in CMV(-) and CMV(+). Cell types with significant frequency differences between CMV(-) and CMV(+) groups are marked with an asterisk (*; P-adjusted [Benjamini-Hochberg] < 0.05). *Right:* Select CD8^+^ T cell frequencies within PBMCs quantified *via* scRNA-seq in Discovery Cohort: in CMV(-) (n=13) and CMV(+) (n=14). **d,** CD8^+^ TEMRA cell frequencies within PBMCs quantified by flow cytometry in Validation Cohort: CMV(-) (n=23) and CMV(+) (n=40). **e,** Absolute count of CD8^+^ TEMRA cells quantified by flow cytometry in Discovery Cohort: CMV(-) (n=8) and CMV(+) (n=8). **f,** γδ T cell frequencies within PBMCs quantified by flow cytometry in Validation Cohort: CMV(-) (n=23) and CMV(+) (n=40). **g,** GZMK^+^ CD8^+^, GZMB^+^ γδ, and CD8^+^ TEMRA T cell frequencies within PBMCs quantified *via* scRNA-seq in Cohort 3: CMV(+) (n=42) and CMV(-) (n=50). **h,** *Left:* UMAP representing 4 subsets from CD4^+^ T cells (n= 55,927). Treg, CD4^+^ T-regulatory; EM, effector memory; TEMRA, terminally differentiated effector memory. *Right:* UMAPs of CD4^+^ T cells showing expression of select marker genes. **i,** UMAP color coded for CMV serostatus. **j,** *Left:* CD4^+^ T cell subset frequencies in CMV(-) and CMV(+). Cell types with significant frequency differences between CMV(-) and CMV(+) groups are marked with an asterisk (*; P-adjusted [Benjamini-Hochberg] < 0.05). *Middle:* Naïve CD4^+^ T cell frequencies within PBMCs. *Right:* CD4^+^ TEMRA cell frequencies within PBMCs. All panels represent scRNA-seq data from Discovery Cohort: CMV(-) (n=13) and CMV(+) (n=14). **k,** CD4^+^ TEMRA frequencies within PBMCs quantified by flow cytometry in Validation Cohort: CMV(-) (n=23) and CMV(+) (n=40). **l,** Absolute count of CD4^+^ TEMRAs quantified by flow cytometry in Discovery Cohort: CMV(-) (n=8) and CMV(+) (n=8). **m,** CD4^+^ TEMRA T cell frequencies within PBMCs quantified *via* scRNA-seq in Cohort 3: CMV(+) (n=42) and CMV(-) (n=50). Box plots display the median and IQR (25–75%), with whiskers representing the upper and lower quartiles ±1.5× IQR. Each dot represents an individual sample. Mann-Whitney-U test (two-sided) was used to compare cell counts between CMV(-) and CMV(+) (**c, d, e, f, g, j, k. l, m**). All *P*-values are shown after adjusting for multiple hypothesis testing using Benjamini-Hochberg at false discovery rate (FDR) < 0.05.

CD8+ TEMRA cells exhibited a 9.5-fold relative expansion in CMV(+) compared CMV(-) adults (7.2% vs 0.8% of PBMCs; p<0.001; Fig. 2c), which was confirmed by flow cytometry in Cohort 2 (p<0.001; Fig. 2d and Extended Data Fig. 2c). Absolute CD8^+^ TEMRA counts supported these results (p=0.001; Fig. 2e). γδ T cells, which recognize antigens independently of MHC presentation^43^, were also more frequent in CMV(+) adults, particularly the GZMB^+^ γδ subset which showed a 5.8-fold increase (3.7% vs 0.6%; p<0.001; Fig. 2c and Extended Data Fig. 2b). Increase in γδ T cell frequency was confirmed in Cohort 2 by flow cytometry (p=0.05; Fig. 2f and Extended Data Fig. 2c). GZMK^+^ CD8^+^ T cells were also more frequent in CMV(+) adults (6% vs 1.6% of PBMCs; p<0.001; Fig 2c and Extended Data Fig. 2b).

To assess whether these CMV-associated alterations were specific to older adults, we re-analyzed publicly available scRNA-seq datasets from younger adults (Cohort 3; n=47, 25-35 years old; n=45, 55-65 years old; Extended Data Fig. 2d) with known CMV serostatus (CMV(+) n=42; CMV(-) n=50)^32^. CD8^+^ TEMRA T, GZMB^+^ γδ T, and GZMK^+^ CD8^+^ T cell frequency was also higher in CMV(+) younger adults (p<0.001; Fig. 2g and Extended Data Fig. 2e).

In-depth analyses of each CD8^+^ subset revealed very few differentially expressed genes (DEGs; FDR<0.05) between CMV(+) and CMV(-) older individuals (Supplementary Table 2). To further explore cell-intrinsic differences, we calculated single-cell scores for cytotoxicity and lymphoid innateness—a gradient that evaluates transcriptional similarity of T cells to innate lymphocytes (Supplementary Table 3)^44,45^. These scores were largely comparable between CMV(+) and CMV(-) donors across all subsets, except for GZMK^+^ CD8^+^ T cells that displayed an increased cytotoxicity score for CMV(+) individuals (Extended Data Fig. 2f, g). Together, CMV seropositivity is associated with substantial compositional remodeling of the CD8^+^ T cell compartment with minor cell-intrinsic alterations in older adults.

### Frequency of CD4^+^ TEMRA T cells is higher in CMV(+) older adults

CD4⁺ T cells (55,927) (Fig. 2h and Extended Data Fig. 3a) were clustered into four subsets: (i) naïve (*CCR7^+^*); (ii) T-regulatory cells (Tregs; *FOXP3*^+^, *TIGIT*^+^), (iii) effector memory (EM; *S100A4*^+^, *GZMB*^-^), and (iv) TEMRA cells (*GZMB*^+^, *GNLY^+^*). CMV(+) individuals exhibited a lower frequency of naïve CD4^+^ T cells (p=0.04) and higher frequency of CD4^+^ TEMRA cells (p=<0.001; Fig. 2j and Extended Data Fig. 3b). CD4^+^ TEMRA cells were 6.4-fold higher in CMV(+) individuals (2.15% vs 0.34% of PBMCs), confirmed by flow cytometry in Cohort 2 (p=0.009; Figure 2k and Extended Data Fig. 2c). Absolute counts of CD4^+^ TEMRA cells were also higher in CMV(+) (p=0.08; Fig. 2l). Cohort 3 displayed similar relative expansion of CD4^+^ TEMRA cells (p<0.001; Fig. 2m and Extended Data Fig. 3c). DEG analysis yielded few (n=3) significant genes (Supplementary Table 2), with *CCL5* upregulated in Tregs from CMV(+) older adults.

Further clustering of the CD4^+^ memory compartment (EM; n=19,691; Extended Data Fig. 3d) revealed five memory subsets (i) T helper 1 (Th1; *IFNG-AS1*^+^, *GZMK*^+^), (ii) T helper 17 (Th17; *RORC*^+^, *CCR6*^+^, *CTSH*^+^), (iii) T follicular helper-like (Tfh-like; *CXCR5*^+^) (iv) T helper 2 (Th2; *GATA3*^+^), and (v) T helper 10 (Th10; *CCR10*^+^)^46^. None of these memory subsets remodeled with CMV seropositivity in our cohort (Extended Data Fig. 3e,f) or in Cohort 3 (Extended Data Fig 3g,h).

Thus, CMV seropositivity in older adults is associated with higher frequency of CD4^+^ TEMRA cells without alterations of other memory CD4^+^ T cell subsets.

### Frequency of atypical B cells (ABCs) is higher in CMV(+) older adults

B cells (n=12,573; Fig. 3a and Extended Data Fig. 4a) clustered into five subsets: (i) naïve (*IL4R*^+^, *IGHM*^+^), (ii) transitional (*CD9*^+^, *MME*^+^), (iii) ABC (*CD27*^-^, *ITGAX*/CD11c^+^, *TBX21*/Tbet^+^), (iv) memory (*CD27*^+^, *IGHG1*^+^, *IGHA1*^+^), and (v) plasmablasts (PB; *JCHAIN*^+^, *MZB1*^+^). CMV(+) individuals had higher frequencies of ABCs, with a 3.4-fold increase (0.27% vs 0.9%; p=0.01; Fig. 3b,c and Extended Data Fig.4b). This increase in frequency was also observed in 55–65-year-old CMV(+) individuals from Cohort 3 (p=0.04; Fig. 3d and Extended Data Fig. 4c). No DEGs were found between CMV(+) and CMV(-) older adults in any B cell subset (Supplementary Table 2).

**Fig. 3.**
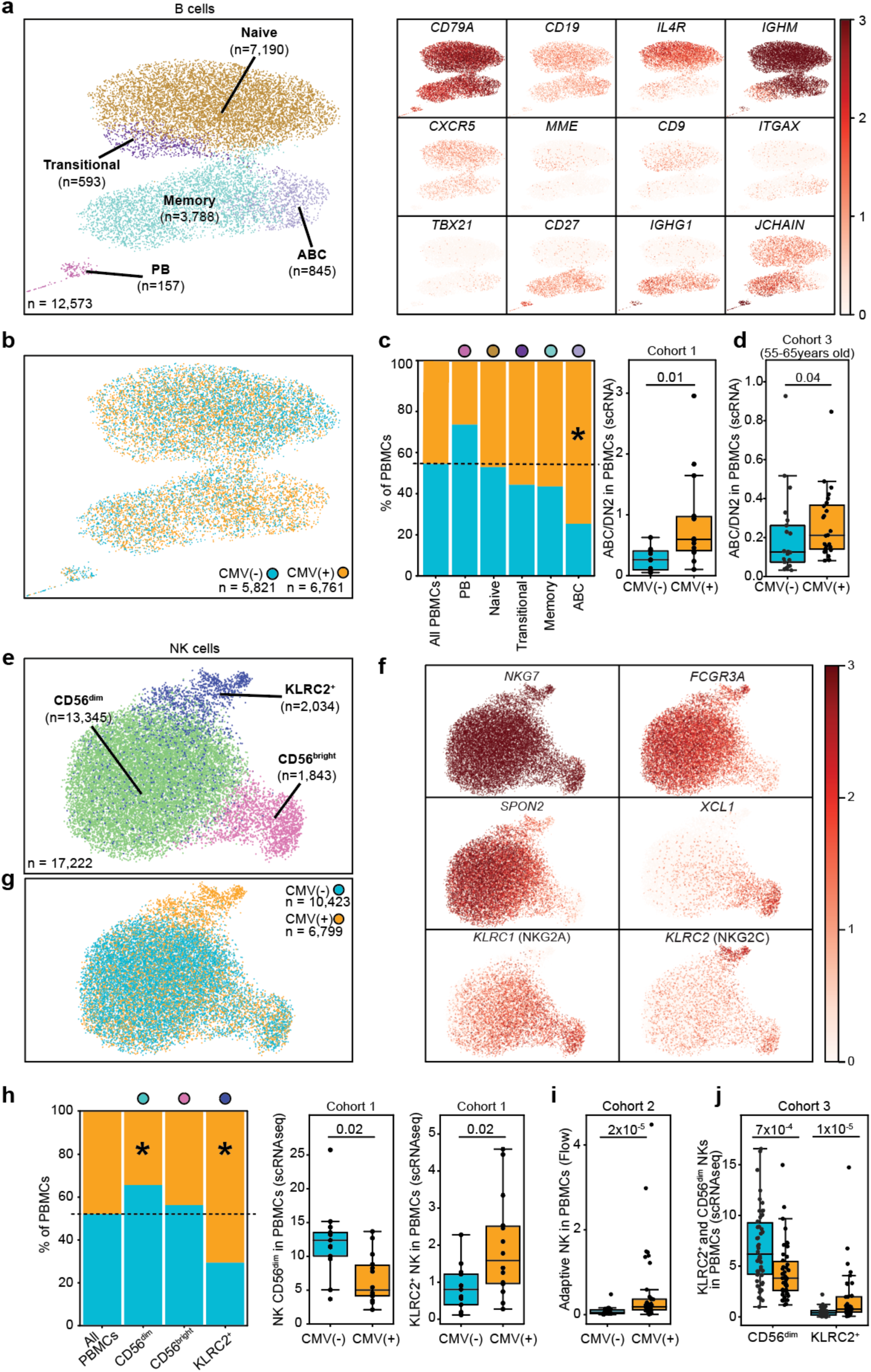
ABC and adaptive NK cells were expanded while CD56^dim^NK cells were reduced in CMV(+) older adults. **a,** *Left:* UMAP representing 5 B cell subsets (n= 12,573). PB, Plasmablast; ABC, atypical B cell. *Right:* UMAPs of B cells showing expression of select marker genes. **b,** UMAP color coded for CMV serostatus. **c,** *Left:* B cell frequencies in CMV(-) and CMV(+). Cell types with significant frequency differences between CMV(-) and CMV(+) groups are marked with an asterisk (*; P-adjusted [Benjamini-Hochberg] < 0.05). *Right:* ABC frequencies within PBMCs All panels represent scRNA-seq data from Discovery Cohort: CMV(-) (n=13) and CMV(+) (n=14). **d,** ABC frequencies within PBMCs quantified *via* scRNA-seq in Cohort 3 (55-65 age group): CMV(+) (n=24) and CMV(-) (n=20). **e,** UMAP representing 3 NK cell subsets (n= 17,222). **f,** UMAPs of NK cells showing expression of select marker genes. **g,** UMAP color coded for CMV serostatus. **h,** *Left:* NK cell frequencies in CMV(-) and CMV(+). Cell types with significant frequency differences between CMV(-) and CMV(+) groups are marked with an asterisk (*; P-adjusted [Benjamini-Hochberg] < 0.05). *Middle:* NK CD56^dim^ frequencies within PBMCs. *Right:* KLRC2^+^ NK cell frequencies within PBMCs. All panels represent scRNA-seq data from Discovery Cohort: CMV(-) (n=13) and CMV(+) (n=14). **i,** Adaptive NK frequencies within PBMCs quantified by flow cytometry in Validation Cohort: CMV(-) (n=23) and CMV(+) (n=40). **j,** NK CD56^dim^ and KLRC2^+^ NK cell frequencies quantified *via* scRNA-seq in Cohort 3: CMV(+) (n=42) and CMV(-) (n=50). Box plots display the median and IQR (25–75%), with whiskers representing the upper and lower quartiles ±1.5× IQR. Each dot represents an individual sample. For Mann-Whitney-U test (two-sided) was used to compare cell counts between CMV(-) and CMV(+) (**c**, **d, h, i, j**). All *P*-values are shown after adjusting for multiple hypothesis testing using Benjamini-Hochberg at false discovery rate (FDR) < 0.05.

Therefore, CMV seropositivity is linked with more frequent ABCs without affecting other B cell subsets.

### Frequency of CD56^dim^NK cells is lower, whereas frequency of NKG2C⁺ NK cells is higher in CMV(+) older adults

NK cells (n=17,222) (Fig. 3e,f) clustered into three subsets: (i) CD56^dim^ NK cells (*SPON2^+^*, *FCGR3A*^+^, *GZMB^+^)*, (ii) CD56^bright^ NK cells (*NCAM1*/CD56 *^+^*, *GZMK^+^*, *XCL1^+^)*, and (iii) KLRC2*^+^* NK cells (*KLRC2^+^*/NKG2C), which closely resembled adaptive NK cells (Extended Data Fig. 4d)^47,48^.

CD56^dim^ NK cell frequency was lower in CMV(+) individuals in both our older adult cohort (6.6% vs 12.2% of PBMCs; p=0.02; Fig. 3g,h) and Cohort 3 (p<0.001; Fig. 3j and Extended Data Fig. 4e). Conversely, KLRC2^+^ NK cell frequency was higher by 2.3-fold in CMV(+) older adults, (2% vs. 0.8%; p=0.011, Fig. 3h), which was confirmed by flow cytometry in Cohort 2 (p<0.001; Fig. 3i and Extended Data Fig. 4f). KLRC2^+^ NK cells were also more frequent in CMV(+) individuals within Cohort 3 (p<0.001; Fig. 3j and Extended Data Fig. 4e). No DEGs were found between CMV(+) and CMV(-) older adults in NK subsets (Supplementary Table 2).

### Monocyte and dendritic cell frequencies are comparable between CMV(+) and CMV(-) older adults

Monocytes (n=30,132) (Fig. 4a and Extended Data Fig. 5a) clustered into four subsets: (i) CD14^+^ monocytes (*CD14^+^*, *LYZ^+^*, *S100A8^+^*, *S100A9^+^*), (ii) ISG-high CD14^+^ monocytes (*CD14^+^*, *IFI44L^+^*, *MX1^+^*), (iii) intermediate monocytes (*FCGR3A*^+^, *CD14*^+^, *HLA*-*DRB1*^+^), and (iv) CD16^+^ monocytes (*FCGR3A*^+^, *FCER1G*^+^). No differences in monocyte subset frequencies or DEGs were observed between CMV(+) and CMV(-) individuals (Fig. 4b, Extended Data Fig. 5b, and Supplementary Table 2). Monocyte subset was also not altered with CMV in Cohort 3 (Extended Data Fig. 5c).

**Fig. 4.**
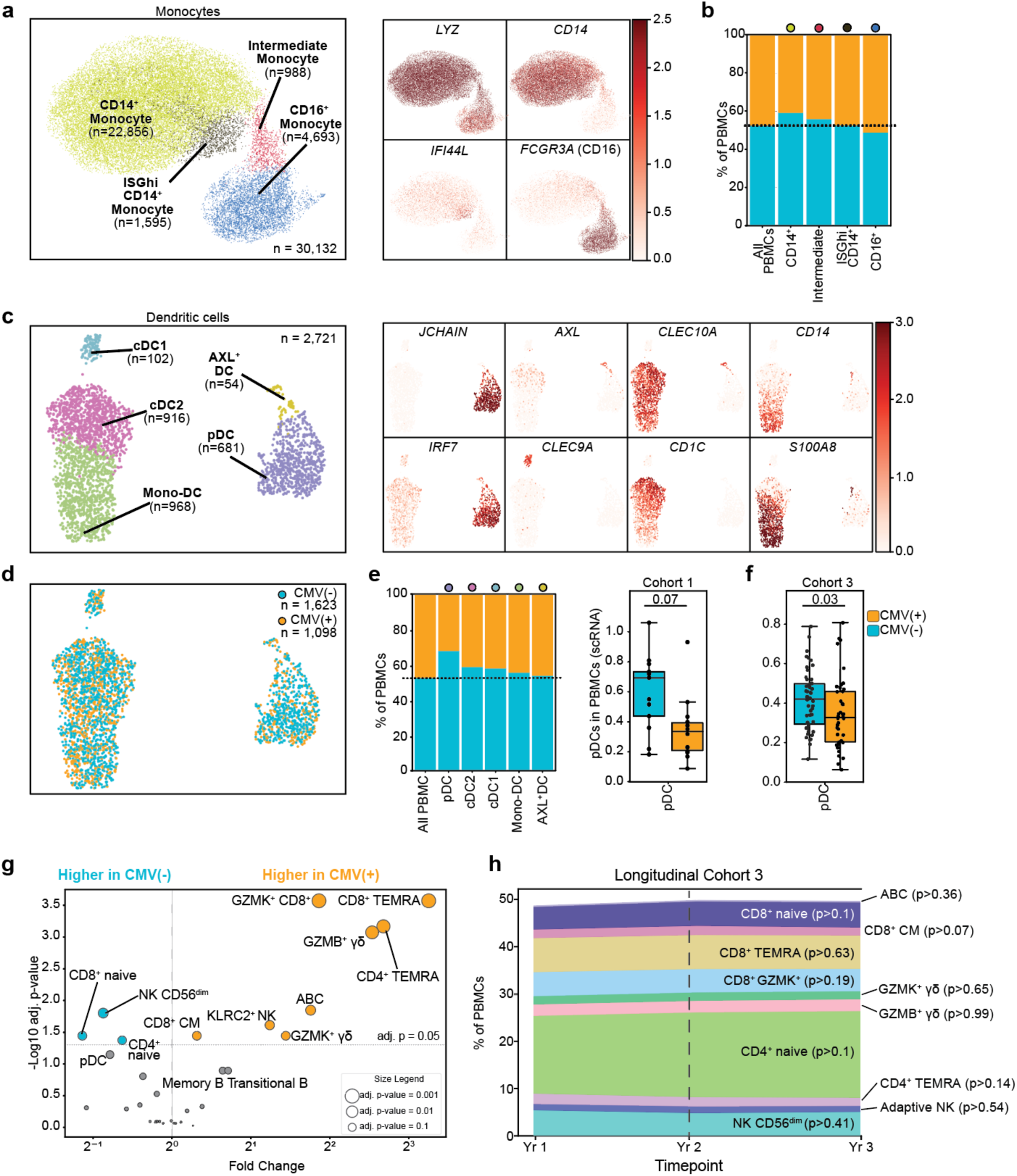
Monocytes and dendritic cells were comparable between CMV(+) and CMV(-) older adults. **a,** *Left:* UMAP representing 4 monocyte subsets (n= 30,132). ISGhi, interferon stimulated gene-high expressing subset. *Right:* UMAPs of monocytes showing expression of select marker genes. **b,** Monocyte frequencies in CMV(-) and CMV(+). **c,** *Left:* UMAP representing 5 DC subsets (n= 2,721). DC, dendritic cell; Mono-DC, monocyte-derived dendritic cell; cDC1, conventional type 1 dendritic cells; cDC2, Conventional type 2 dendritic cells; pDC, plasmacytoid dendritic cell. *Right:* UMAPs of DCs showing expression of select marker genes. **d,** UMAP color coded for CMV serostatus. **e,** *Left:* DC frequencies in CMV(-) and CMV(+). *Right:* pDC frequencies within PBMCs. All panels represent scRNA-seq data from Discovery Cohort: CMV(-) (n=13) and CMV(+) (n=14). **f,** pDC frequencies quantified *via* scRNA-seq in Cohort 3: CMV(+) (n=42) and CMV(-) (n=50). **g,** Summary volcano plot showing PBMC compositional alterations associated with CMV serostatus in Discovery Cohort. Size of bubbles represent p-adjusted value (-Log scale). **h,** Average frequencies of CMV-associated cell types over a three-year period in CMV(+) individuals from Cohort 3 (n=22). For **e,** and **f,** box plots display the median and IQR (25–75%), with whiskers representing the upper and lower quartiles ±1.5× IQR. Each dot represents an individual sample. Mann-Whitney-U test (two-sided) was used to compare cell counts (**e, f, h**). All *P*-values are shown after adjusting for multiple hypothesis testing using Benjamini-Hochberg at false discovery rate (FDR) < 0.05.

Dendritic cells (DCs) (Fig. 4c) clustered into five subsets (i) monocyte-derived dendritic cells (Mono-DC; *S100A8*^+^, *CD14*^+^, *CLEC10A*^+^), (ii) type-2 classical dendritic cells (cDC2; *CLEC10A*^+^, *CD1C*^+^), (iii) type-1 classical dendritic cells (cDC1; *CLEC9A*^+^), (iv) AXL^+^ dendritic cells (AXL^+^ DC; *AXL*^+^), and (v) plasmacytoid DCs (pDCs; *IRF7*^+^, *JCHAIN*^+^). Among these subsets, pDCs showed a trend toward lower frequency in CMV(+) individuals (0.35% vs 0.6% of PBMCs) (p=0.07; Fig. 4, d,e and Extended Data Fig. 5d). A modest but significant reduction in pDC frequency was also observed in Cohort 3 (0.35% vs 0.41%; p=0.03; Fig. 4f and Extended Data Fig. 5e). No DEGs were found between CMV(+) and CMV(-) older adults in DC subsets (Supplementary Table 2).

Overall, our older adult cohorts revealed pronounced CMV-associated remodeling of lymphoid immune cell subsets, with minimal alterations in myeloid cells (Fig. 4g and Supplementary Table 4). To estimate the contribution of clinical factors to lymphocyte frequency variance in Cohort 1, we conducted Principal Variance Component Analysis^49^ (for exact procedure see supplementary materials under “Data and code availability”), which identified CMV serostatus as the dominant contributor (43%), followed by BMI (11%) (Extended Data Fig. 5g). Finally, leveraging longitudinal samples from Cohort 3, revealed that CMV-associated immune signature remains stable over three years (Fig. 4h and Extended Data Fig. 5f), underscoring the persistent nature of latent CMV-driven immune remodeling.

### CMVerify: a novel tool that accurately infers CMV serostatus from PBMC scRNA-seq data

To enable analysis of public single-cell RNA-seq studies lacking CMV serostatus information, we developed a machine learning model: CMVerify. This tool leverages a Random Forest classification model that uses PBMC subtype frequencies as input (i.e., data features) to predict latent CMV serostatus (i.e., classification output) (Fig. 5a). CMVerify was trained on Cohort 3 (n=92 total; n=42 CMV(+); n=50 CMV(-); Extended Data Fig. 2c) using leave-one-out cross-validation, achieving 96.7% training accuracy. Top predictive features included CD4^+^ TEMRA, CD8^+^ TEMRA, KLRC2^+^ NK cells, and GZMB^+^ γδ T cells (Extended Data Fig. 6a,b), consistent with our CMV-associated immune signature. We first validated CMVerify in our Discovery Cohort, where it correctly predicted CMV serostatus in 26 out of 27 individuals (Fig. 5b and Extended Data Fig. 6c). We then further tested CMVerify on two independent older adult cohorts with different chemistries using available CMV IgG and PBMC scRNA-seq data: Cohort 4 (n=19, age 62-90) profiled using 10X 3’ scRNAseq, Cohort 5: (n=20, age 64-88) profiled using 10X 5’ scRNAseq. Across these datasets, CMVerify achieved 97% average accuracy, 98% positive predictive value (PPV), and 96% negative predictive value (NPV) (Fig. 5b, Extended Data Fig. 6c).

**Fig. 5.**
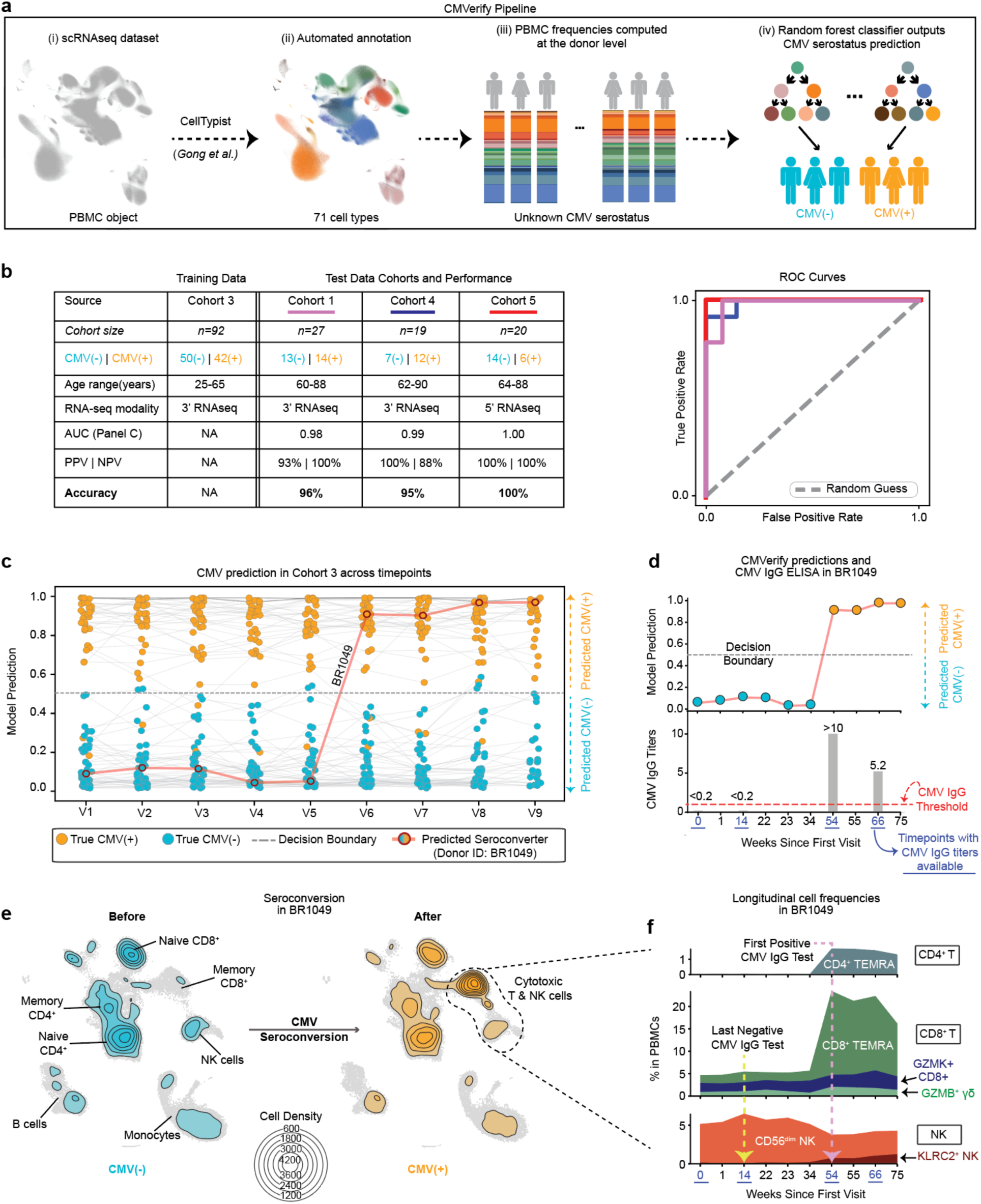
*CMVerify*—a novel tool that accurately predicts CMV serostatus from scRNA-seq data. **a,** Schematic representation of *CMVerify* Pipeline: (i) *CMVerify* accepts a PBMC scRNA-seq dataset as input, (ii) applies CellTypist to automatically annotate the dataset, (iii) computes PBMC frequencies at the donor level, and (iv) uses a random forest classifier to predict CMV serostatus. **b,** *Left*: Table summarizing *CMVerify’s* training dataset and performance in three independent test cohorts spanning different age groups and scRNA-seq modalities. PPV, positive predictive value; NPV, negative predictive value; *Right*: Receiver Operating Characteristic (ROC) curves showing performance of *CMVerify* in the independent cohorts. **c,** *CMVerify* prediction results for the Cohort 3 longitudinal data. Y-axis represents model predicted probability, each bubble is a unique sample corresponding with a donor timepoint. Samples from the same donor are connected via gray lines. Based on the Decision Boundary (y = 0.5), all bubbles below the Decision Boundary (y < 0.5) represent prediction of CMV(-), vs. bubbles above the Decision boundary (y > 0.5) represent prediction of CMV(+). Bubbles are color coded for true CMV serostatus (determined via CMV IgG ELISA). Red line connects model predicted probabilities for the seroconverter (donor ID: BR1049; n = 1) across all timepoints (n = 9). **d,** *Top*: *CMVerify* prediction probabilities across all timepoints in donor BR1049 identified in panel **c**. *Bottom*: CMV IgG titers for donor BR1049. Underlined timepoints (in blue) indicate presence of CMV IgG data at those timepoints. The red dotted line indicates CMV IgG titer threshold above which a sample is considered CMV(+). **e**, Contour UMAP plots show PBMC densities before and after CMV seroconversion in donor BR1049. The UMAP labeled ‘Before’ includes timepoints at weeks 14, 22, 23, and 34, while the UMAP labeled ‘After’ includes weeks 54, 55, 66, and 75. The Cell Density legend indicates the number of cells within each contour level shown in the UMAPs. **f**, CMV-driven changes in Cytotoxic T and NK cell frequencies over time in donor BR1049.

To evaluate longitudinal performance, CMVerify was applied to 77 individuals from Cohort 3, each with PBMC scRNA-seq data across 9 timepoints. The model maintained 97% accuracy, correctly classifying 670 of 693 longitudinal samples (Fig. 5c and Extended Data Fig 6d). Misclassifications primarily stemmed from two donors with unusually low CD4⁺ TEMRA frequencies (Extended Data Fig 6e). Remarkably, in a donor who seroconverted over the study (35-year-old male, ID: BR1049), CMVerify successfully tracked the transition, shifting its prediction from CMV(-) to CMV(+) in parallel with rising CMV IgG titers (Fig. 5c,d). This transition was accompanied by rapid (within months) and dramatic changes in PBMC composition, including a 90.99-fold increase in CD4⁺ TEMRAs, 8.74-fold increase in CD8⁺ TEMRAs, and 3.37-fold increase in KLRC2⁺ NK cells (Fig. 5e and Extended Data Fig. 6f). These analyses established that CMVerify is an effective tool that can be used to infer CMV serostatus from single cell datasets in the absence of serological data.

### CMV-specific TCR clones are expanded in GZMK^+^ CD8^+^ T and Th1 cells

To study how CMV-seropositivity alters T cell clonal diversity, we profiled PBMCs of 20 older adults (Cohort 5) using 10X 5’ scRNA-seq with paired TCR sequencing (Fig. 5b and Fig. 6a). We recovered 128,924 high-quality TCRα–TCRβ pairs, averaging 6,446 T cells per donor (Fig. 6a), with comparable cell counts between CMV(-) and CMV(+) groups (Extended Data Fig. 7a). These 128,924 T cells represented 103,575 clones, where a clone was defined as a unique TCRα–TCRβ pair at the nucleotide level. We first quantified overall TCR repertoire diversity using the exponent of Shannon’s entropy (e^H^) across all memory CD4⁺ and CD8⁺ T cells for each donor (using the Alakazam package^50^, see Methods). CMV(+) donors had reduced overall TCR diversity compared to CMV(-) donors (Fig. 6b and Supplementary Table 5), in line with previous observations^30^. Subset-specific analysis revealed that GZMK⁺ CD8⁺ T cells exhibited reduced TCR clonal diversity in CMV(+) individuals, in addition to CD4^+^/CD8^+^ TEMRA cells (Fig. 6c). Surprisingly, Th1 cells also had lower TCR diversity in CMV(+) individuals (Fig. 6c and Extended Data Fig. 7b,c; see Methods), despite showing no change in frequency (Extended Data Fig. 3f).

**Fig. 6.**
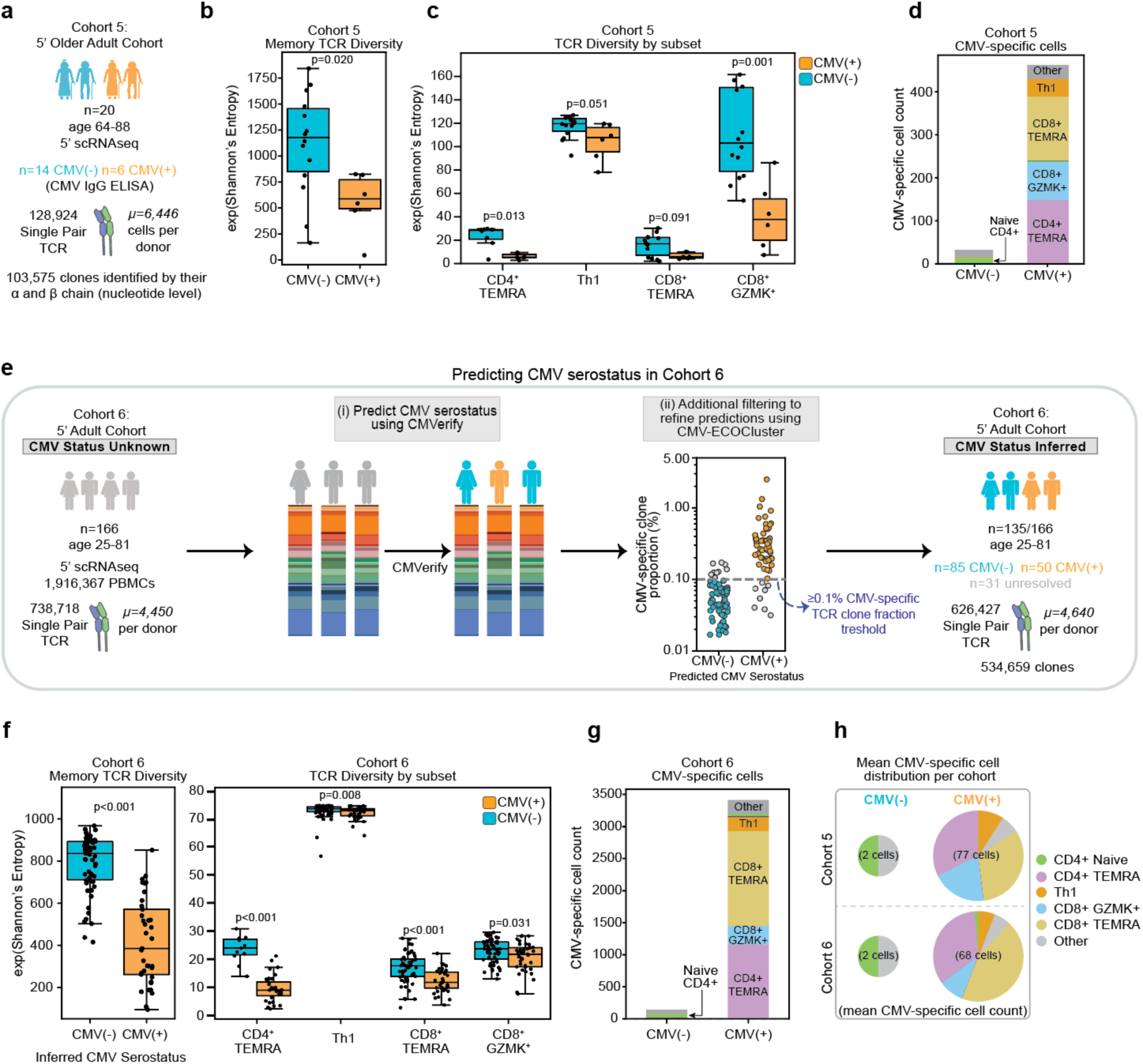
Putative CMV-specific TCR clones are expanded in GZMK^+^ CD8^+^ and Th1 T cells. **a**, Cohort 2 included *n=14* CMV(-) and *n=6* CMV(+) older adults profiled using 5’ scRNAseq with paired scTCR-seq. **b**, Cohort 2 TCR diversity in memory T cells. **c,** Cohort 2 TCR diversity in CD4^+^/CD8^+^ TEMRA, Th1, and CD8^+^ GZMK^+^ T subsets. **d**, Cohort 2 putative CMV-specific cell counts identified by exact matching with the CMV ECOcluster (see Methods). **e**, Schematic representation of method to infer CMV serostatus in Cohort 3 (*n=166* donors with CMV status unknown). (i) predict CMV serostatus using CMVerify, (ii) additional filtering to refine CMVerify predictions based on the fraction of putative CMV-specific clones: <0.1% in predicted CMV(–) individuals and ≥0.1% in predicted CMV(+) individuals. The gray line (y = 0.1%) indicates the cutoff threshold (derived from Cohort 2 with ELISA CMV IgG; Extended Data Fig. 7d). In total, CMV-serostatus was confidently inferred for 135/166 individuals in Cohort 3. **f**, *Left:* Cohort 3 TCR diversity in memory T cells. *Right:* Cohort 3 TCR diversity in CD4^+^/CD8^+^ TEMRA, Th1, and CD8^+^ GZMK^+^ T subsets. **g**, Cohort 3 putative CMV-specific cell counts identified by exact matching with the CMV ECOcluster. **h**, Distribution of CMV-specific TCR by cell type in CMV(-) and CMV(+) individuals, grouped by cohort.. The size of each pie chart represents the average number of CMV-specific cells (log scale). The mean CMV-specific cell count per group and cohort is indicated in parentheses. For **b, c,** and **f,** box plots display the median and IQR (25–75%), with whiskers representing the upper and lower quartiles ±1.5× IQR. Each dot represents an individual sample. Mann-Whitney-U test (two-sided) was used to compare differences between groups. TCR diversity was calculated using Shannon’s entropy (e^H^). For **c**, due to the small sample size in the CMV(+) group (6/20), diversity calculations within cell types were not FDR-corrected. For **f**, *P*-values shown are after adjusting for multiple hypothesis testing using Benjamini-Hochberg at false discovery rate (FDR) < 0.05.

To identify putative CMV-specific TCRs, we leveraged the CMV ECOcluster, a recently curated database of 26,106 TCRβ sequences enriched in over 12,000 CMV(+) individuals^51^. A subset of the ECOcluster TCRβ chains were previously experimentally validated as responding CMV peptides via the VDJdb database^52^ (primarily pp65 and IE1; see Methods; Supplementary Table 5). By matching exact V–J pairings and CDR3 amino acid sequences between the CMV ECOcluster and TCR-seq data from our cohort, we identified 493 putative CMV-specific cells (Fig. 6d) from 115 clones (see Methods). In CMV(-) donors, we identified a small population (32 cells from 30 clones; Extended Data Fig. 7d) of naïve CD4^+^ T cells (based on their scRNA-seq transcriptional profile; Extended Data Fig. 7e,f), consistent with stochastic V(D)J recombination in naïve T cells^53,54^ and absence of antigen-driven differentiation (Fig. 6d). In contrast, CMV(+) donors had clonally expanded CMV-specific T cells (461 cells from 85 clones; Fig. 6d and Extended Data Fig. 7a,e). As expected, the highest frequencies of putative CMV-specific cells were detected within the CD4⁺ TEMRA (mean clonal expansion: 11.5x) and CD8⁺ TEMRA (10.6x) subsets (Fig. 6d and Extended Data Fig. 7g). CMV-specific TCRs were also expanded within the GZMK⁺ CD8⁺ T cells (4.4x; Fig. 6d and Extended Data Fig. 7g), consistent with their increased frequency (Fig. 2c) and reduced TCR diversity (Fig. 6c). Finally, the Th1 helper subset contained a modest expansion of CMV-specific clones (1.5x) (Fig. 6d and Extended Data Fig. 7g), in line with its decreased TCR diversity (Fig. 6c).

To validate our observations in an independent cohort, we obtained a publicly available PBMC scRNA-seq dataset with paired scTCR-seq from 166 donors (aged 25–81 years, Cohort 6)^2^ (Fig. 6e). Since CMV serostatus was not reported, we first used CMVerify to obtain CMV serostatus predictions. To increase confidence in our downstream analysis, we independently confirmed and refined CMVerify predictions with the CMV ECOcluster (see Methods) to confidently resolve CMV serostatus of 135 donors: 85 predicted CMV(-) and 50 predicted CMV(+) (Fig. 6e, Extended Data 7h, and Supplementary Table 6). We identified 626,427 high-quality TCRα–β pairs (averaging 4,640 per donor) and repeated our TCR diversity analysis in this cohort. Reproducing our findings in Cohort 5, CMV(+) individuals in Cohort 6 also had reduced overall TCR diversity and CMV-specific clonal expansion, particularly within CD4⁺/CD8⁺ TEMRA, GZMK⁺ CD8⁺ T, and Th1 subsets (Fig. 6f,g, Extended Data 7i, and Supplemental Table 6). In total, we identified 141 CMV-specific cells in CMV(-) individuals (mostly within the naïve CD4^+^ T subset) and 3,410 CMV-specific cells in CMV(+) individuals (Fig. 6g). Importantly, in CMV(+) individuals, these CMV-specific cells were detected in CD4⁺/CD8⁺ TEMRA, GZMK⁺ CD8⁺ T, and Th1 subsets, as observed in our Cohort 5 (Fig. 6g). Overall, across both cohorts, the average CMV(-) individual possessed one CD4⁺ naïve CMV-specific cell (Fig. 6h). In contrast, CMV(+) individuals had an average of 73 CMV-specific cells, of which 71% were CD4^+^/CD8⁺ TEMRA, 15% were GZMK⁺ CD8⁺ T and 8% were Th1 cells (Fig. 6h).

By analyzing single-cell RNA-seq paired with TCR-seq data across two cohorts, we showed that within the CD8⁺ lineage, there is CMV clonal expansion of GZMK⁺ CD8⁺ T cells in addition to CD8⁺ TEMRA cells^55^. Within the CD4^+^ lineage, CMV-specific clones were detected in Th1 cells along with CD4^+^ TEMRA cells^33^.

## Discussion

This is the most comprehensive study of CMV-associated changes in cellular composition and clonal diversity, integrating data from six human cohorts spanning different age groups, including two newly built older adult cohorts. We confirmed known associations in CD4⁺/CD8⁺ TEMRA T cells^24,26,32,33,55,56^, γδ T cells^34^, and KLRC2⁺ (adaptive) NK cells^35,36^. Importantly, we expand these immune signatures by showing that: (i) GZMK⁺ CD8⁺ T cells are increased in frequency and display CMV-specific clonal expansion, (ii) CD4⁺ Th1 helper cells exhibit CMV-specific clonal expansion, and (iii) atypical B cells are increased in frequency and CD56^dim^ NK cells are reduced in CMV(+) individuals across age groups.

GZMK⁺ CD8⁺ T cells have been traditionally linked to aging and inflammaging^2,2,57^. Here, we show that these cells are increased in frequency with latent CMV, exhibit reduced TCR diversity, and harbor expanded CMV-specific clones in both younger and older CMV(+) adults, supporting a direct role for CMV in their expansion. Similar GZMK signatures were also observed in human tissues with latent CMV^58^. GZMK is a tryptase-like protease recently shown to activate complement cascade, initiating a lymphocyte-driven inflammatory response^59^. These data suggest that inflammation may also contribute to the immune control of viral latency, in addition to the well-defined cytotoxic responses (e.g., CD4^+^/CD8^+^ TEMRA and adaptive NK cells). How inflammatory and cytotoxic pathways and cell types interact to sustain CMV control, however, remains to be elucidated.

Our TCR analyses across two independent cohorts uncovered Th1 cells as the only CD4^+^ T helper subset harboring CMV-specific clones expanded in CMV(+) individuals, albeit with smaller clone sizes. These findings suggest that CMV-driven clonal expansion occurs within the CD4^+^ T helper subset but appears more tightly regulated as compared to CD4^+^ TEMRA cells. Future studies will be essential to understand how Th1 cells contribute to latent CMV responses, e.g., by helping cytotoxic lymphocyte proliferation^60^ and/or stimulating CMV-specific antibody formation via ABCs (Fig. 3c,d).

CMV serostatus is often missing in scRNA-seq datasets, despite its known influence on immune phenotypes. To retrospectively analyze such datasets, we developed CMVerify—a machine learning classifier that predicts CMV serostatus from single-cell transcriptomic data with high accuracy. We provide CMVerify as an open-source tool for the research community. Prior tools can leverage cytometry^61^ or TCR-seq^62^ data to predict CMV serostatus, while CMVerify is the first model to do so using scRNA-seq data. Despite being trained on a younger adult cohort, CMVerify accurately predicted CMV status across three independent older adult cohorts, highlighting the robust and age-independent imprint of CMV on immune cell compositions. Furthermore, CMVerify successfully captured a seroconversion event, highlighting rapid (within months) PBMC compositional shifts linked to acute/latent CMV. CMVerify was instrumental in reproducing our CMV-specific TCR analyses and novel observations in GZMK⁺ CD8⁺ T and Th1 cells by allowing us to infer CMV status in a public dataset^2^.

Our comprehensive multi-modal study revealed novel CMV-associated changes in cellular composition and TCR diversity, including the clonal expansion of GZMK⁺ CD8⁺ T and Th1 cells in CMV(+) individuals. Future studies should aim to understand how cytotoxic (CD4⁺/CD8⁺ TEMRA T cells), inflammatory (GZMK⁺ CD8⁺ T cells), and T helper (Th1 cells) cells orchestrate effective control of latent CMV in healthy adults.

### Limitations

Our study has several limitations. First, the exciting results observed in CMV seroconversion are unfortunately based on a single case (*n* = 1). Out of ∼250 individuals followed over two years, only one seroconversion event was captured. Therefore, substantially larger cohorts will be needed to capture additional seroconversion events and establish a causal relationship between CMV infection and the rapid remodeling of circulating immune cell populations. Second, the CMV ECOcluster was derived from enrichment analyses and has not yet been fully experimentally validated. While the ECOcluster is large and highly specific (with some TCRs mapping to the experimentally validated VDJdb database; see Methods), further experimental validation of these TCR sequences will be instrumental. Despite these limitations, our study provides valuable and novel insights into the systemic effects of latent CMV infection across multiple cohorts and age groups.

## Materials and Methods

### Cohort Descriptions

**Cohort 1** (n=36) comprised individuals recruited from two separate studies:

1. UConn Center on Aging Bacterial Pneumonia Vaccine Study: Twenty healthy older adults aged 60 years and older were recruited at the University of Connecticut Center on Aging during the 2017 and 2018 seasons (IRB number: 16-071J; R01AG052608; R01AI142086). Participants did not have confounding treatments or conditions that would influence immune function and composition (e.g., immunotherapy, immunosuppressive disorders, and prednisone in doses greater than 10 mg/day)^63^.
2. Immune Response to High-Dose vs. Standard Dose Influenza Vaccine at the University of Connecticut Health Center (UCHC): Sixteen healthy older adults aged 65 years and older were recruited at UCHC and Health Sciences North Research Institute (HSNRI) in 2017 to study resilience to influenza infections. Exclusion criteria for these participants included known immunosuppressive disorders or medications, including prednisone in doses greater than 10 mg/day.

The CMV(+) and CMV(-) groups in Cohort 1 had comparable distributions in age, sex, body mass index (BMI), frailty index, medication use, and cancer history (see Supplementary Table 1).

**Cohort 2** (n=63): This publicly available dataset^40,41^ was profiled via flow cytometry. It consisted of healthy older adults (66-92 years old). The CMV(+) and CMV(-) groups in Cohort 2 had comparable distributions in age, sex, BMI, frailty index, and cancer history (see Supplementary Table 1).

**Cohort 3** (n=96): This publicly available dataset^32^ was profiled using 10X 3’ scRNA-seq and included longitudinal samples from n=49 healthy young adults (25-35 years old) and n=47 healthy older adults (55-65 years old).

**Cohort 4** (n=19): This publicly available dataset^64^ was profiled using 10X 3’ scRNA-seq and included n=95 samples, of which we used n=19 samples (62-90 years old) with known CMV serology (ELISA) data.

**Cohort 5** (n=20): Healthy older adults (64-88 years old) were recruited at the University of Connecticut Center on Aging during the 2022-2023 seasons (IRB number: 2022-Ceded-002) and profiled using 10X 5’ scRNA-seq with paired scTCR-seq. Participants did not have confounding treatments or conditions that would influence immune function and composition (e.g., immunotherapy, immunosuppressive disorders, and prednisone in doses greater than 10 mg/day).

The CMV(+) and CMV(-) groups in Cohort 5 had comparable distributions in age, sex, BMI, ongoing cancer diagnosis, and frailty index (see Supplementary Table 1).

**Cohort 6** (n=166): This publicly available dataset^2^ was profiled using 10X 5’ scRNA-seq with paired scTCR-seq and included longitudinal samples from n=166 healthy adults (25-81 years old).

### Summary of Cohorts Used in the Study

**Table 1.** Summary of cohorts used in this study. For each cohort, the table lists the source, profiling modality of PBMCs, age distribution, CMV seropositivity distribution, and the specific purpose of use in this study.

| Cohort name | Originating Study | Modality | Age range | CMV seropositivity | Purpose of use |
| --- | --- | --- | --- | --- | --- |
| Cohort 1 | Grabauskas et al. | a) 10X 3' scRNA-seq<br>b) Flow cytometry | 60-88 | CMV(-): 22<br>CMV(+): 21 | a) Derive CMV-linked immune signatures<br>b) Test CMVerify |
| Cohort 2 | Verschoor et al.<br>Picard et al. | Flow cytometry | 66-92 | CMV(-): 23<br>CMV(+): 40 | Validate CMV-linked immune signatures |
| Cohort 3 | Gong et al. | 10X 3' scRNA-seq | 26-66 | CMV(-): 52<br>CMV(+): 44 | a) Validate CMV-linked immune signatures<br>b) Train CMVerify |
| Cohort 4 | Nehar-Belaid et al. | 10X 3' scRNA-seq | 62-90 | CMV(-): 7<br>CMV(+): 12 | Test CMVerify |
| Cohort 5 | Grabauskas et al. | a) 10X 5' scRNA-seq<br>b) scTCR-seq | 64-88 | CMV(-): 14<br>CMV(+): 6 | a) Test CMVerify<br>b) CMV-linked TCR analysis |
| Cohort 6 | Terekhova et al. | a) 10X 5' scRNA-seq<br>b) scTCR-seq | 25-80 | NA | Validate CMV-linked TCR findings |

### Sample Collection in Cohort 1 and Cohort 5

In Cohort 1 and Cohort 5, peripheral blood mononuclear cells (PBMCs) were isolated by Lymphoprep (StemCell Technologies) gradient centrifugation within 1h after collection of blood samples by Lymphoprep (StemCell Technologies) gradient centrifugation whereas serum was isolated from the blood collected in red top collection tubes (BD Vacutainer). The isolated PBMCs were resuspended in a cryoprotective solution containing 90% FBS and 10% DMSO, then transferred to liquid nitrogen for long-term storage whereas serum was stored at -80°C.

### Cytomegalovirus Serostatus in Cohort 1 and Cohort 5

CMV serostatus was determined by ELISA using the CMV IgG ELISA kits (Aviva Systems Biology and Genesis Diagnostics Ltd; see Supplementary Table 1). Serum samples were thawed at room temperature, diluted, and analyzed according to manufacturer’s recommendations. Calculation of the results was done based on the appropriate controls included in the kit.

### 10X 3’ Single-Cell Library Preparation and Sequencing in Cohort 1

In Cohort 1, we generated a total of 27 baseline PBMC scRNA-seq samples from older adults as part of two separate studies. All of these samples were generated at the Jackson Laboratory Single Cell Biology Lab using the same pipeline. Briefly, cells were washed and resuspended in PBS containing 0.04% BSA and immediately processed as follows. Cell viability was assessed on a Countess FL II automated cell counter (Thermofisher) using Trypan Blue staining, and up to 12,000 cells from each suspension were loaded onto one lane of a 10x Genomics Chip G. Single cell capture, barcoding, and library preparation were performed using the 10x Genomics Chromium Controller platform version 3.1 NEXTGEM chemistry and according to the manufacturer’s protocol (CG000315, 10x Genomics). cDNA and libraries were assessed for quality via Tapestation 4200 (Agilent) and Qubit fluorometer (ThermoFisher), quantified by KAPA qPCR, and sequenced on a NovaSeq 6000 (Illumina) S4 v1.5 200 flow cell using a 28-10-10-90 asymmetric read configuration targeting a read depth of 50,000 per cell.

### 10X 5’ Single-Cell Library Preparation and Sequencing in Cohort 5

In Cohort 5, we generated a total of 20 baseline PBMC scRNA-seq samples from older adults. All samples were generated at the Jackson Laboratory Single Cell Biology Lab using the same pipeline. After dissociation, cells were washed and suspended in PBS containing 0.04% BSA and immediately processed as follows. Cell viability was assessed on a LUNA FX7 automated cell counter (Logos Biosystems) via AO/PI staining, and up to 29,000 cells from each suspension were loaded onto one lane of a 10x Genomics Chromium GEM-X Single Cell 5’ Chip v3. Single cell capture, barcoding and library preparation were performed using the 10x Genomics Chromium X platform^65^ 5’ version 3 GEM-X chemistry and according to the manufacturer’s protocol (CG000733, revA). From each individual sample, cDNA was derived and used for the construction of a gene expression and a TCR (VDJ) library. cDNA and libraries were checked for quality by Tapestation 4200 (Agilent) and Qubit Fluorometer (ThermoFisher) and quantified by KAPA qPCR. Libraries were sequenced on an Illumina NovaSeq X+ 25B 300 cycle flow cell, with a 28-10-10-90 (R1-I1-I2-R2) asymmetric read configuration, targeting 20,000 barcoded cells with an average sequencing depth of 50,000 reads per cell and 15,000 reads per cell for the gene expression and TCR library.

### scRNA-seq preprocessing in Cohort 1, Cohort 3, and Cohort 5

In <u>Cohort 1 and Cohort 5</u>, Illumina base call files for all libraries were converted to FASTQs using bcl2fastq v2.20.0.422 (Illumina, CA, USA) and FASTQ files associated with baseline gene expression libraries were aligned to the GRCh38 reference assembly with v32 annotations from GENCODE (10x Genomics GRCh38 reference 2020-A) using the version 6.1.1 Cell Ranger count pipeline (10x Genomics, CA, USA), as previously described^63^. The resulting gene-cell expression matrices, were then corrected for ambient RNA contamination using SoupX^66^. The corrected gene-cell expression matrices were then used for downstream analysis. We used the Scrublet^67^ package in Python to estimate the doublet score for each sample with an estimated doublet set to 0.1. Doublets were manually confirmed to be doublets via expression of PBMC lineage markers before being removed from downstream analyses. Cells with (i) genes that were not detected in at least 3 cells, (ii) mitochondrial reads greater than 25% (20% in Cohort 5), and (iii) fewer than 200 features were excluded. Filtered gene expression matrices were merged and processed using the Scanpy toolkit (version 1.9.1)^68^. Read counts were log normalized to the total number of reads in each individual cell to count-per-million (CPM) using the “normalized_total” and “log1p” functions. Genes with the highest normalized dispersion were identified using the “highly_variable_genes” function. Gene expression matrices were scaled using “scale” function and Principal Component Analysis (PCA) was performed using the “pca” function. To correct batch effects across samples at the PBMC level, we applied Harmony^69^ in Cohort 1 and Cohort 5. To correct batch effects across samples at the lineage level, we applied Harmony in Cohort 1 and SCVI-Tools^70^ in Cohort 5. For the baseline samples, 100 Principal Components (PCs) were used to calculate nearest neighbors via the “neighbors” function which were used to perform uniform manifold approximation and projection (UMAP) *via* the “umap” function. Initial clusters were determined using the “leiden” function (resolution=0.8) for all PBMCs. Multiple rounds of marker identification, cell type annotation, and manual inspection and doublet removal, were performed to annotate PBMC subsets in Cohort 1 and Cohort 5.

In <u>Cohort 3</u>, our compositional analysis focused on the Flu Year 1 Day 0 baseline time point (Fig. 2g,m, Extended Data Fig. 2c, Fig.3d,j, and Fig.4f), which included n=1,539,411 cells. This dataset was filtered—by the original authors—for doublets using Scrublet and further refined by removing cells with more than 5,000 genes or fewer than 200 genes, and mitochondrial threshold over 10%. The only additional threshold we imposed for automated quality control was to remove genes detected in less than 3 cells, which left 32,069 genes detected. Cells were clustered and annotated using the same process we applied to Cohort 1. In Figure 2h we also include annotations following the above procedure for Flu Year 2 Day 0 (Yr 2 V1) and Year 3 Stand-Alone (Yr 3 V1) timepoints. Finally, in Figure 5e for the seroconverter we annotated cells from all available timepoints.

### *CMVerify* model training

Baseline samples (Flu Year 1 Day 0) from Cohort 3 (n=92)^32^ were used for training *CMVerify*. The input to our machine learning pipeline was donor-level PBMC frequencies of the 72 cell types, labeled with the Allen Institute for Immunology Level 3 (High Resolution, i.e., AIFI_L3) CellTypist model^32,71^. PBMC frequencies were scaled using “StandardScaler” as provided by the scikit-learn package^72^ to transform frequencies to have zero mean and unit variance. A random forest classifier was used for model training. Optimal parameters were selected using grid search and leave-one-out cross-validation over all possible combinations of the following hyperparameters:

Optimal parameters (indicated in bold in Table 2) yielded the best cross validation score of 0.967.

**Table 2:** Parameters tested and selected (in bold) for CMVerify training.

| <u>*Parameter*</u> | <u>*Grid Search Options (selected)*</u> |
| --- | --- |
| number of estimators | 50, **100**, 200 |
| max depth | **None**, 10, 20, 30 |
| minimum samples to split internal node | 2, **5**, 10 |
| minimum samples required at a leaf | 1, 2, **4** |
| number of features to consider when looking for best split | **square root (number of features)**, log<sub>2</sub> (number of features) |
| sampling with replacement | True, **False** |

Feature importances were calculated based on the mean decrease in impurity where impurity is a measure of how mixed a node in a tree is with respect to target labels. Features (cell type frequencies) that best reduce the impurity and more effectively split the CMV(+) and CMV(-) samples will have the highest importance (Extended Data Fig. 6a, Table 3). Fourteen features were selected by *CMVerify* based on feature importance scores (Table 3). To evaluate how the Cell Typist labeling corresponds with our Discovery Cohort cell type labels (see Methods:scRNA-seq data analyses), we mapped the top five *CMVerify* selected Cell Typist features to our own cell type annotations (Extended Data Fig. 6b).

**Table 3:** Fourteen features were selected by *CMVerify* during training procedure, listed here in descending order of importance (for predicting CMV serostatus).

| *Cell Type* | *Importance* |
| --- | --- |
| KLRF1- GZMB+ CD27- memory CD4 T cell (CD4<sup>+</sup> TEMRA) | 0.351594 |
| KLRF1- GZMB+ CD27- EM CD8 T cell (CD8<sup>+</sup> TEMRA) | 0.162693 |
| Adaptive NK cell (KLRC2<sup>+</sup> NK) | 0.137851 |
| KLRF1- effector Vd1 gdT (GZMB<sup>+</sup> $\gamma\delta$) | 0.092140 |
| KLRF1+ effector Vd1 gdT (GZMB<sup>+</sup> $\gamma\delta$) | 0.090094 |
| Proliferating NK cell | 0.033280 |
| GZMK- CD27+ EM CD8 T cell | 0.026493 |
| GZMK+ CD56dim NK cell | 0.025931 |
| GZMK- CD56dim NK cell | 0.024149 |
| KLRF1+ GZMB+ CD27- EM CD8 T cell | 0.018953 |
| ISG+ CD56dim NK cell | 0.010061 |
| GZMK+ memory CD4 Treg | 0.009613 |
| pDC | 0.009209 |
| CD8 MAIT | 0.007939 |

### TCR analyses in Cohort 5

Samples from Cohort 5 were processed using the same methods as described above (see Methods: scRNA-seq preprocessing in Cohort 1, Cohort 3, and Cohort 5). Cells were annotated based on their scRNA-seq transcriptional profiles, and paired TCR-seq data was analyzed following the Scirpy pipeline^73^. Before quality control, we obtained 159,626 cells which had paired TCR-seq data, of which 14% had more than one pair of TCRs, and 5% had orphan TCRs. We excluded those 29,605 cells that did not have a single pair from the analysis. We also excluded cells annotated as γδ T cells, leaving 128,924 CD4^+^/CD8^+^ T cells with a single pair TCR after quality control (6,446 cells on average from each donor, see Fig. 6a). Cell counts were comparable between CMV(-) and CMV(+) donors (see Extended Data Fig. 7a); although CMV(+) donors had more CD4^+^ TEMRA (Supplementary Table 5). We computed the distance between TCR using the identity metric at the nucleotide level, defining the clonotypes based on both receptor arms’ primary chain^73^ (Table 4).

**Table 4:**
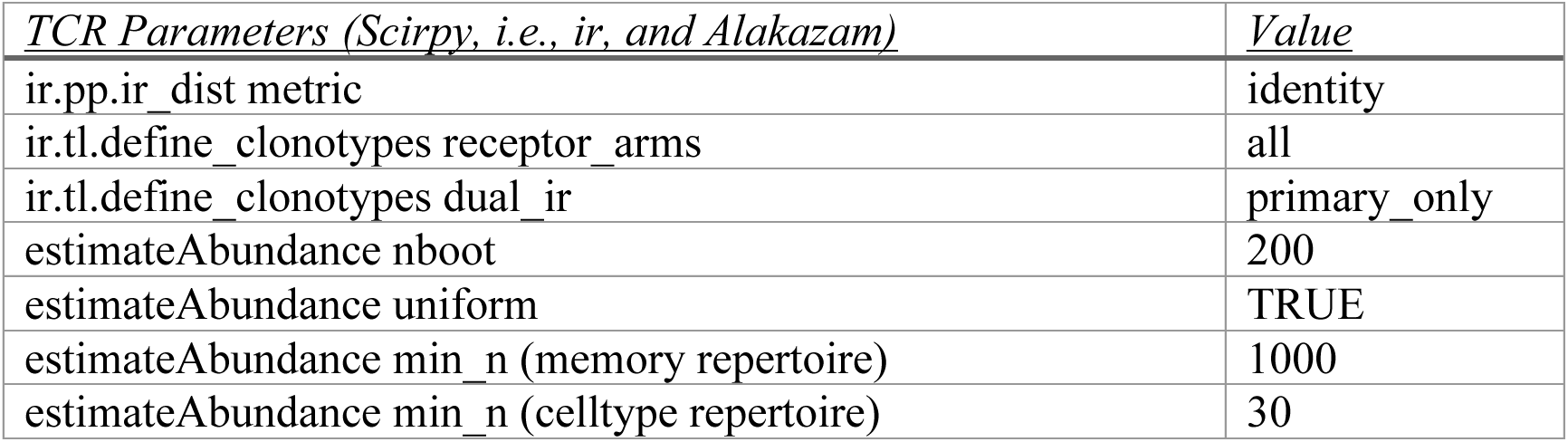
Parameters for TCR analysis via Scirpy and Alakazam to enable reproducibility. Column one lists the function name and then specific parameters with parameter value in column two.

To estimate TCR diversity per cell type, we computed the Hill Diversity Index ^q^D^74^ where q=1, (i.e., the exponent of Shannon Entropy, e^H^), using the ‘estimateAbundance’ method from the Alakazam package^50^. We used the uniform sampling parameter to control for differences in cell counts between donors, i.e., all samples were ‘downsampled’ to the minimum across samples during the procedure which included 200 bootstrap replicates (Table 4). To measure CMV-specific clonal expansion in the memory compartment (see Fig. 6b), we excluded naïve CD4^+^, naïve CD8^+^, and naïve Treg cells. To identify subset specific differences in repertoire diversity between CMV(-) and CMV(+) donors, we again used the ‘estimateAbundance’ function (same parameters as above), with the additional restriction that each donor must have at least 30 cells within the subset to be included in the comparison (see Fig. 6c and Extended Data Fig.7c).

### Putative CMV-specific TCR identification in Cohort 5 and Cohort 6

To identify putative CMV-specific TCRs, we searched the CMV ECOcluster^51^ (see Methods: CMV ECOcluster) for exact matches with TCR in Cohort 5 and Cohort 6, i.e., identical TCRβ V gene, J gene and CDR3 amino acid sequence. Using Cohort 5 (with known CMV IgG serology via ELISA), we established a CMV-specific clone fraction cutoff that divided our CMV(-) and CMV(+) donors: all CMV(-) donors had <0.1% of their total clones matched as CMV-specific (see Extended Data Fig. 7d). CMV-specific clone fraction for each individual was calculated as follows:

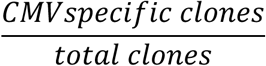

### CMV status prediction in Cohort 6

We applied CMVerify to predict CMV status in Cohort 6 using the 1,916,367 PBMCs from 166 donors who each had samples across 1-3 timepoints. We applied CMVerify for each sample and timepoint. In total, 94 donors were consistently predicted CMV(-) and 65 were consistently predicted as CMV(+) across all timepoints (see Extended Data Fig. 7h). We excluded seven donors with inconsistent CMVerify predictions as unresolved. To further refine our predictions and increase confidence in the downstream analysis, we utilized our established cutoff (based on Cohort 5 where CMV serostatus was determined with CMV IgG ELISA; see Extended Data Fig. 7d and Methods: Putative CMV-specific TCR identification in Cohort 5 and Cohort 6) of <0.1% of total clones matched as CMV-specific. Specifically, we excluded: (i) CMVerify predicted CMV(-) individuals with CMV-specific clone proportion ≥0.1%, and (ii) CMVerify predicted CMV(+) individuals with CMV-specific clone proportion <0.1% (see Fig. 6e). To summarize, we confidently resolved 135/166 donors CMV serostatus in Cohort 6, with 50 individuals predicted CMV(+) and 85 individuals predicted CMV(-) (see Fig. 6e).

### TCR analyses in Cohort 6

Cohort 6 contained 738,718 single pair TCR from donors across the lifespan^2^ (see Fig 6e). We first downloaded their publicly available single-cell RNA- and TCR-seq data, matching the donor metadata using paired barcodes. We followed an identical TCR quality control procedure as described above. Note that we made three minor changes to their scRNA-seq cell annotations to enable better comparison with Cohort 6 annotations: (1) combining CD8^+^ Tem GZMB^+^ and CD8^+^ TEMRA to be called CD8^+^ TEMRA (both were *GZMB*^+^), (2) combining CD4^+^ Terminal effector and CD4^+^ TEMRA to be called CD4^+^ TEMRA (both were GZMB^+^), and (3) combining Th1/Th17 and Th1 to be called Th1 (both were IFNG-AS1^+^, GZMK^+^). Due to low cell counts (compared to Cohort 5), we combined TCR data from all timepoints for each individual in the TCR diversity calculations. Overall cell counts were comparable between predicted CMV(-) and CMV(+) (Supplementary Table 6); although predicted CMV(+) donors had more CD4^+^/CD8^+^ TEMRA, GZMK^+^ CD8^+^, Proliferative CD8^+^, and KLRC2^+^ CD8^+^ (Supplementary Table 6). We computed the Hill Diversity index for each individual’s memory repertoire using 98 donors that met the criteria for inclusion (minimum number of cells 1000, see Table 4). We followed the same procedure to compute subset level repertoire diversity as described above (see Fig 6f and Methods: TCR analyses in Cohort 5). To identify CMV-specific TCRs in Cohort 6, we followed the same procedure as described above (see Methods: Putative CMV-specific TCR identification in Cohort 5 and Cohort 6).

We provide antigen gene information, cell type breakdown, donor metadata, and all alpha and beta chain information at both amino acid and nucleotide level for all CMV-specific clones identified in Cohort 5 and Cohort 6 (see Supplementary Table 5, 6).

### CMV ECOcluster

To identify CMV-specific TCRs, we obtained the publicly available CMV ECOcluster (Exposure Co-Occurrence cluster)^51^ which was previously identified using a multi-stage procedure outlined in brief as follows. First, HLA genotypes were measured from direct sequencing of 4,144 subjects, in combination with bulk TCR (>200,000 T cells sequenced) for each individual, to identify 1,000,000 public TCR statistically associated with HLA^75^. Leveraging the co-ocurrence patterns of HLA-restricted TCRs, in combination with bulk TCR data from 30,674 individuals, a single TCR cluster was identified for each of 7 common exposures: EBV, HSV-1, HSV-2, Parvovirus, T. gondii, CMV, and SARS-CoV-2^76^. Finally, the CMV ECOCluster, which comprises 26,106 TCRs, was made publicly available^51^. Note that this approach is based solely on the combination of TCRβ V gene, J gene and CDR3 amino acid sequence.

To validate CMV ECOcluster specificity, we examined whether any CMV ECOcluster TCRβ chains had been previously identified in the VDJdb database^52^, which includes 23,755 experimentally-validated CMV-specific TCRβ chains. Among these, 737 CMV-specific TCRβ chains overlapped with the CMV ECOcluster dataset: pp65 (534 matches), IE1 (111 matches), IE2 (89 matches), and pp50 (3 matches). To estimate background matching, we also quantified overlaps with non-CMV-specific TCRβ chains in VDJdb, identifying 398 matches out of 86,818 non-CMV-specific TCRβ chains. Fisher’s exact test revealed a highly significant enrichment of overlap between putative CMV-specific TCRβ chains from the ECOcluster and experimentally validated CMV-specific TCRβ chains in VDJdb (odds ratio=6.65, p<0.0001). This result indicates that putative CMV-specific sequences were over six times more likely to occur among validated CMV-specific TCRs than expected by chance, supporting the robustness of CMV ECOcluster and its specificity.

To identify putative CMV-specific TCRβ chains in Cohort 5, we processed our TCRβ chains from IMGT format^77^ to enable exact matching (based on identical TCRβ V gene, J gene and CDR3 amino acid sequence) with the CMV ECOcluster.

### Differential expression analyses in Cohort 1

Raw RNA counts were aggregated across cell types using the function “ADPBulk” from adpbulk package (GitHub: noamteyssier, 2021, adpbulk). For each cell type, samples containing fewer than 10 cells were excluded from the differential analysis. Raw counts were first converted to a “DGEList” object, and transformed to normalized matrices using the “calcNormFactors” and “cpm” functions in edgeR^78^. Genes with zero normalized counts across all samples within a given cell type were removed. Differentially expressed gene (DEG) analysis was performed using generalized linear models, employing the default normalization method based on the trimmed mean of M-values (TMM) and controlling for batch, sex and site of sample collection. P-values were adjusted using Benjamini–Hochberg correction for multiple comparisons and genes with an adjusted P-value (FDR) < 0.05 were selected as DEGs.

### Batch correction for gene expression in Cohort 1

In Cohort 1, our scRNA-seq samples included two scRNA-seq libraries (previously published, n=11; and unpublished, n=16) that were sequenced separately. Batch effects were corrected as follows: For each cell type of interest, raw RNA counts were aggregated at the donor level (pseudobulk) as described above. Next, pseudobulk matrix values were adjusted for batch using pyComBat^79^. Finally, the mean expression of genes of interest were calculated at the donor level using batch-corrected pseudobulk matrix.

### Gene scoring in Cohort 1

Cytotoxicity score for each sample was calculated by computing the mean expression of “Cytotoxic lymphocytes” (M9.1) gene set in CD8^+^ T cells ^44^ (n=29 genes, Supplementary Table 3). Innateness score was calculated by computing the mean expression of innateness-related genes in CD8^+^ T cells^45^ (n=80 genes, Supplementary Table 3).

### Flow cytometry data generation and analyses in Cohort 1

The absolute counts of the major cell populations in PBMCs from CMV(+) and CMV(-) individuals in Cohort 1 were labeled and analyzed as follows. Fluorescent-labeled antibody cocktails for cell-surface staining panels were pre-mixed in BD Horizon Brilliant Stain Buffer (BD Biosciences) 10 minutes before staining. Antibody cocktails were added over 100 μL aliquots of anticoagulated whole blood in a 5 ml FACS tube within 60 minutes of blood collection. Samples were incubated for 15 minutes at room temperature then lysed and fixed with 2 ml of 1x FACS lysing solution (BD Biosciences) for 8 minutes at room temperature. The lysed samples were washed twice to remove the unbound antibodies, lysed RBCs, and platelets and finally resuspended in 250 μL of PBS to which 50 μL of count beads suspension (Count Bright Absolute Counting Beads, Thermo Fisher) were added for the detection of absolute cell counts. Cells were stained with fluorochrome-labeled antibodies to the following surface markers: CD3 (clone UCHT1, BioLegend), CD4 (clone RPA-T4, BD Biosciences), CD8 (clone SFC121Thy2D3, Beckman-Coulter), CD19 (clone HIB19, BD Biosciences), CD14 (clone MSE2, BD Biosciences), CD197 (clone 150503, BD Biosciences), and CD45RA (clone HI100, BD Biosciences), and CD127 (clone HIL-7R-M21, BD Biosciences). The stained cells were acquired with LSR Fortessa X-20 (BD) and analyzed with FlowJo software (BD).

### Statistical tests

In Cohort 1, Cohort 2, and Cohort 5, categorical participant characteristics including sex, race, previous cancer diagnosis, and medications used were compared using Fisher’s exact test (two-sided) or the chi-square test, as applicable, using the SciPy package (GitHub: SciPy, 2011).

Across the study, Mann-Whitney U test (two-sided) was performed to compare CMV(+) and CMV(-) groups for continuous variables (e.g., age, BMI, and frailty), cell frequencies and gene set scores (e.g., Cytotoxicity score, Innateness score). Unless stated otherwise in figure legends, p-values in the main and supplementary figures are reported after multiple hypothesis testing correction (i.e. q) by Benjamini-Hochberg’s procedure at FDR < 0.05.

### Ethics Approval for Cohort 1, Cohort 2, and Cohort 5

The study protocols were approved by the Institutional Review Board of the University of Connecticut Health Center (UConn Health IRB #20-130-J-1 or JAX IRB #2019-006-JGM and IRB# - 2022-Ceded-002) and the Health Sciences North Research Ethics Board (#985). Written informed consent was obtained from all participants prior to inclusion in the study.

## Acknowledgments

We gratefully acknowledge the contribution of the Single Cell Biology service, Genome Technologies service, and Cyberinfrastructure high performance computing resources at The Jackson Laboratory (JAX) for assistance with the work described herein. These shared services are supported in part by the JAX Cancer Center (P30 CA034196). We thank Olivia Bart for help with dbGAP data upload. We thank members of the Ucar lab for critical feedback during the progress of the study.

## Author contributions

Conceptualization: JB, GK, DU

Methodology: JB, GK, CPV, DU

Investigation: TG, CPV, LT, AT, DN-B, RM, EP, KES, CCE, HEW, SP, AGS, CK, GY

Visualization: TG, LT, GE

Funding acquisition: JB, GK, CCE, DU

Project administration: TG, DU

Supervision: JB, GK, DU

Writing – original draft: TG, CPV, DU, LT

Writing – review & editing: TG, CPV, DU, JB, GK, LT, AT, DN-B, RM, CK, EP, KES, CCE, HEW, SP, AGS, GY

## Data and code availability

- Processed data and figure code is available on Synapse (under accession code https://doi.org/10.7303/syn68150215). Data are also browsable online (https://latent-cmv-older-adults-400136930153.us-central1.run.app). Raw read data will be made available on dbGAP upon acceptance of the manuscript (under accession code phs003598.v1.p1).

*CMVerify* is available as a Python package: https://pypi.org/project/CMVerify/; open source code hosted at https://github.com/UcarLab/CMVerify.

---

**Extended Data Fig. 1:**
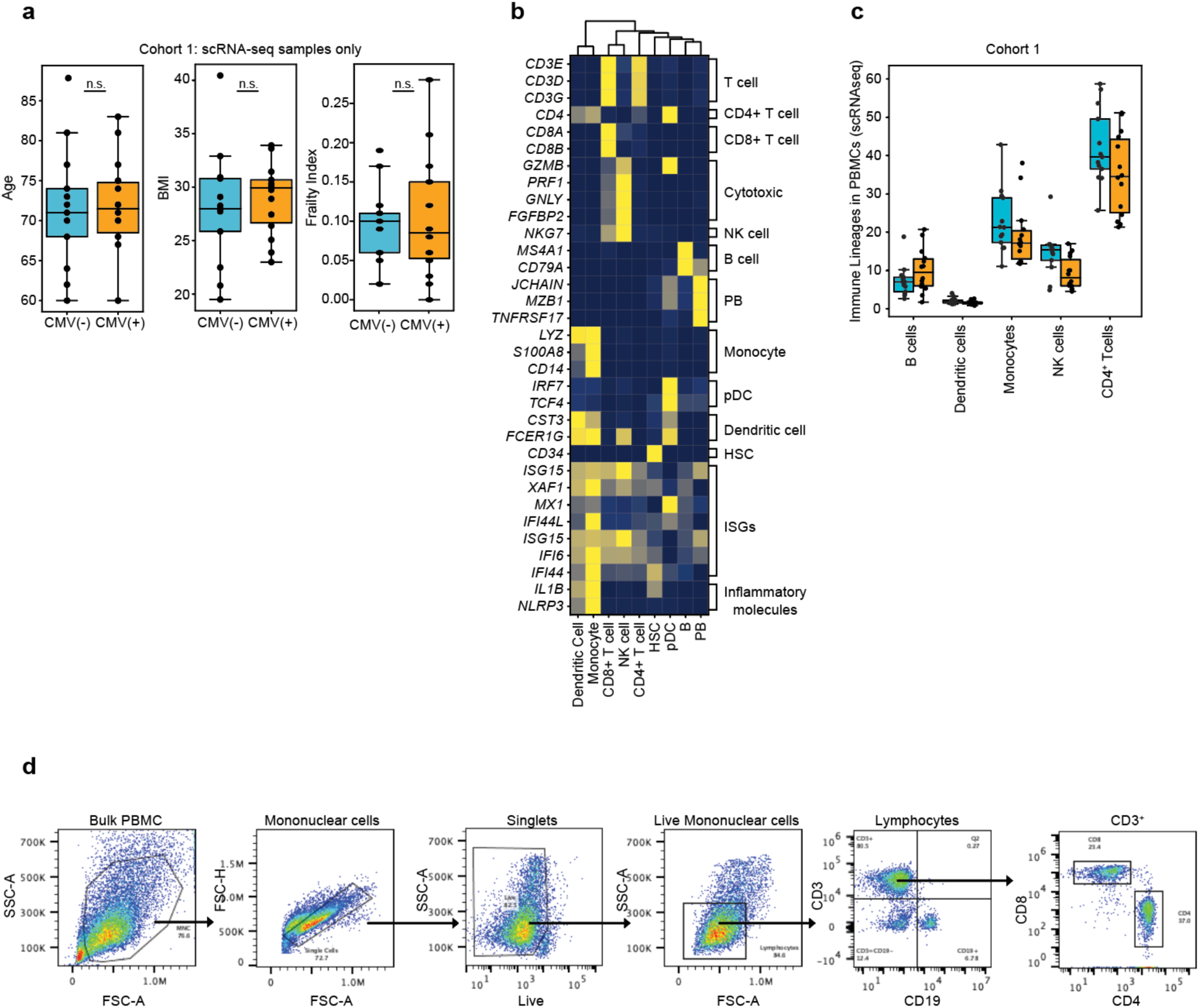
PBMC-level annotations and analysis. **a,** Distribution of age, BMI, and frailty index in Discovery Cohort (scRNA-seq samples only n=27). n.s. indicates not significant. **b,** Scaled expression of select marker genes. **c,** PBMC subset frequency ratios quantified *via* scRNA-seq in CMV(-) (n=13) and CMV(+) (n=14) in Discovery Cohort. **d**, Gating strategy for CD8^+^ and CD4^+^ T cells in Cohort 2 using flow cytometry; FSC, forward scatter; SSC, side scatter. Box plots display the median and IQR (25–75%), with whiskers representing the upper and lower quartiles ±1.5× IQR. Each dot represents an individual sample. For a and c, Mann-Whitney-U test (two-sided) was used to calculate *P*-values between CMV(-) and CMV(+). **c**, none of the measurements were significant after adjusting the *P*-values for multiple hypothesis testing, using Benjamini-Hochberg at false discovery rate (FDR) < 0.05.

**Extended Data Fig. 2:**
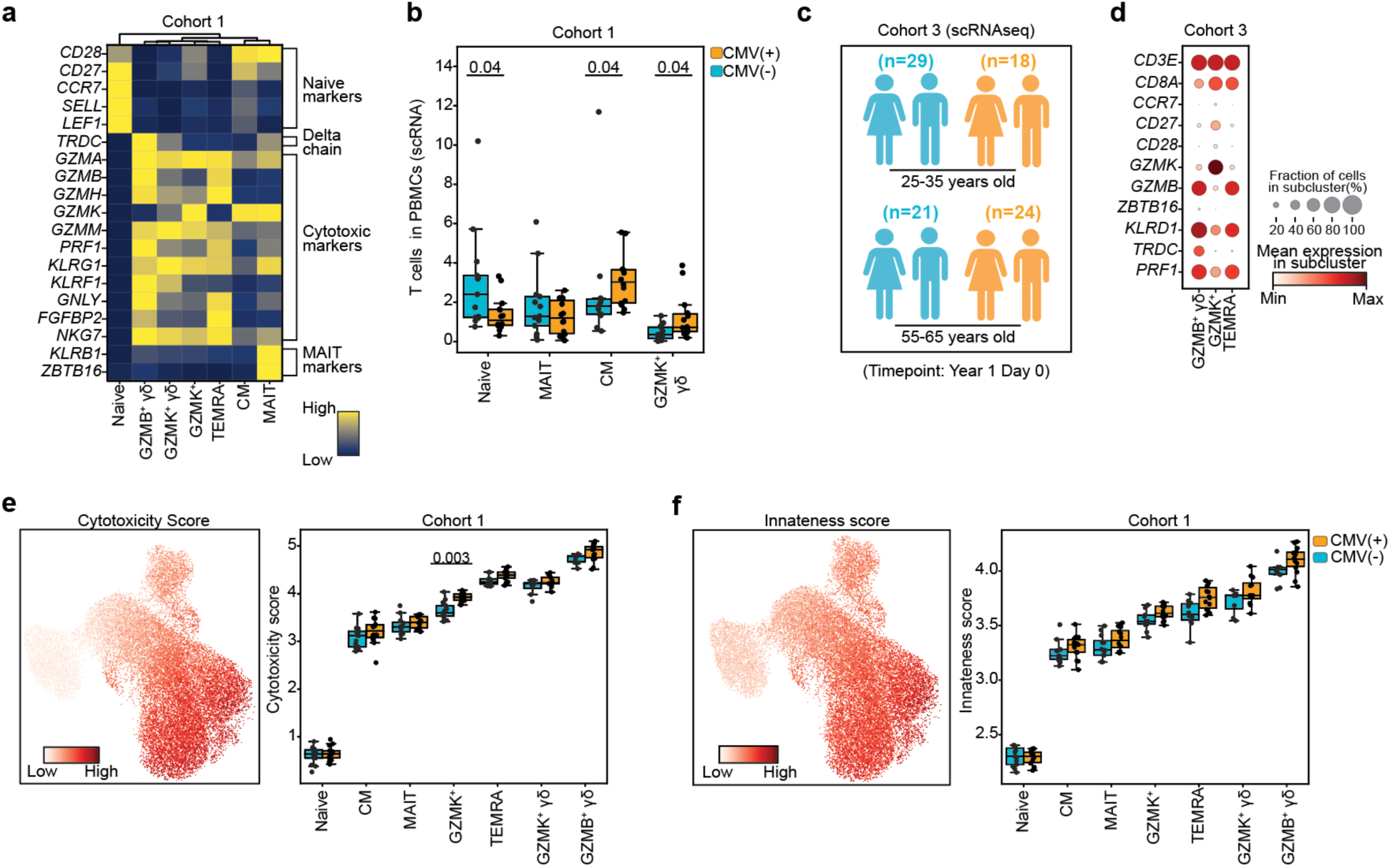
Cytotoxicity and innateness scores for CD8^+^ T cell subsets. **a,** Scaled expression of select marker genes in CD8^+^ T cell subsets. **b,** Select CD8^+^ T cell frequencies within PBMCs quantified *via* scRNA-seq in Discovery Cohort: CMV(-) (n=13) and CMV(+) (n=14). **c**, Gating strategy for CD8^+^/CD4^+^ TEMRA and γδ T cells in Cohort 2 using flow cytometry; FSC, forward scatter; SSC, side scatter. **d,** Schematic representation of Cohort 3 including two age groups (25-35 years old; 55-65 years old). **e,** Scaled expression of select marker for GZMB^+^ γδ T, GZMK^+^ T, and CD8^+^ TEMRA T cells in Cohort 3. **f,** *Left:* UMAP showing Cytotoxicity score in CD8^+^ T cell subsets. *Right:* individual-level cytotoxicity score in CD8^+^ T cells subsets. **g,** *Left:* UMAP showing Innateness score in CD8^+^ T cell subsets. *Right:* individual-level Innateness score in CD8^+^ T cells subsets. Box plots display the median and IQR (25–75%), with whiskers representing the upper and lower quartiles ±1.5× IQR. Each dot represents an individual sample. Mann-Whitney-U test (two-sided) was used to compare cell counts between CMV(-) and CMV(+) (**b, f, g**). All *P*-values are shown after adjusting for multiple hypothesis testing using Benjamini-Hochberg at false discovery rate (FDR) < 0.05.

**Extended Data Fig. 3:**
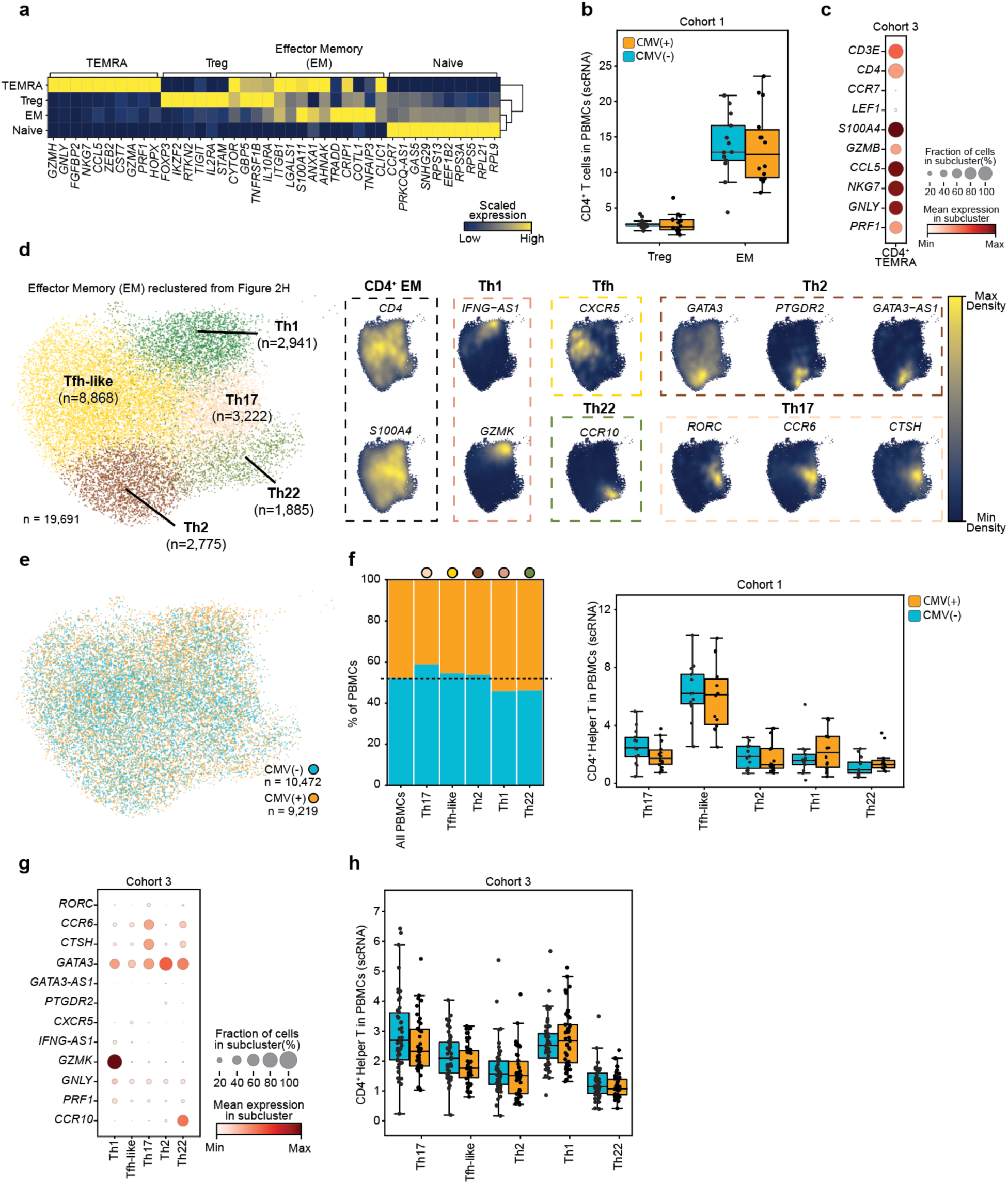
CD4^+^ T helper subsets were comparable between CMV(-) and CMV(+). **a,** Scaled expression of top 10 marker genes in CD4^+^ naïve, CD4^+^ Treg, CD4^+^ EM, and CD4^+^ TEMRA subsets. **b,** CD4^+^ Treg and CD4^+^ TEMRA T cell frequencies within PBMCs quantified *via* scRNA-seq in Discovery Cohort: CMV(-) (n = 13) and CMV(+) (n = 14). **c,** Scaled expression of select marker genes for CD4^+^ TEMRA T cells in Cohort 3. **d,** *Left:* UMAP representing re-clustered CD4^+^ EM compartment and 5 resulting T helper subsets. Tfh-like, T follicular helper-like; Th2, T helper 2; Th22, T helper 22; Th17, T helper 17; Th1. *Right:* Gene density plot showing expression levels of key marker genes, calculated using Nebulosa. Each gene density is grouped according to known T helper marker genes. **e,** UMAP color coded for CMV serostatus. **f,** *Left:* CD4^+^ T helper frequencies in CMV(-) and CMV(+). *Right:* T helper cell frequencies within PBMCs quantified *via* scRNA-seq in Discovery Cohort: CMV(-) (n=13) and CMV(+) (n=14). **g,** Scaled expression of select marker genes for T helper subsets in Cohort 3. **h,** T helper cell frequencies within PBMCs quantified *via* scRNA-seq in Cohort 3: CMV(+) (n=42) and CMV(-) (n=50). Box plots display the median and IQR (25–75%), with whiskers representing the upper and lower quartiles ±1.5× IQR. Each dot represents an individual sample. Mann-Whitney-U test (two-sided) was used to compare cell counts between CMV(-) and CMV(+) (**b**, **e, h,**). None of the measurements were significant after adjusting the *P*-values for multiple hypothesis testing, using Benjamini-Hochberg at false discovery rate (FDR) < 0.05.

**Extended Data Fig. 4:**
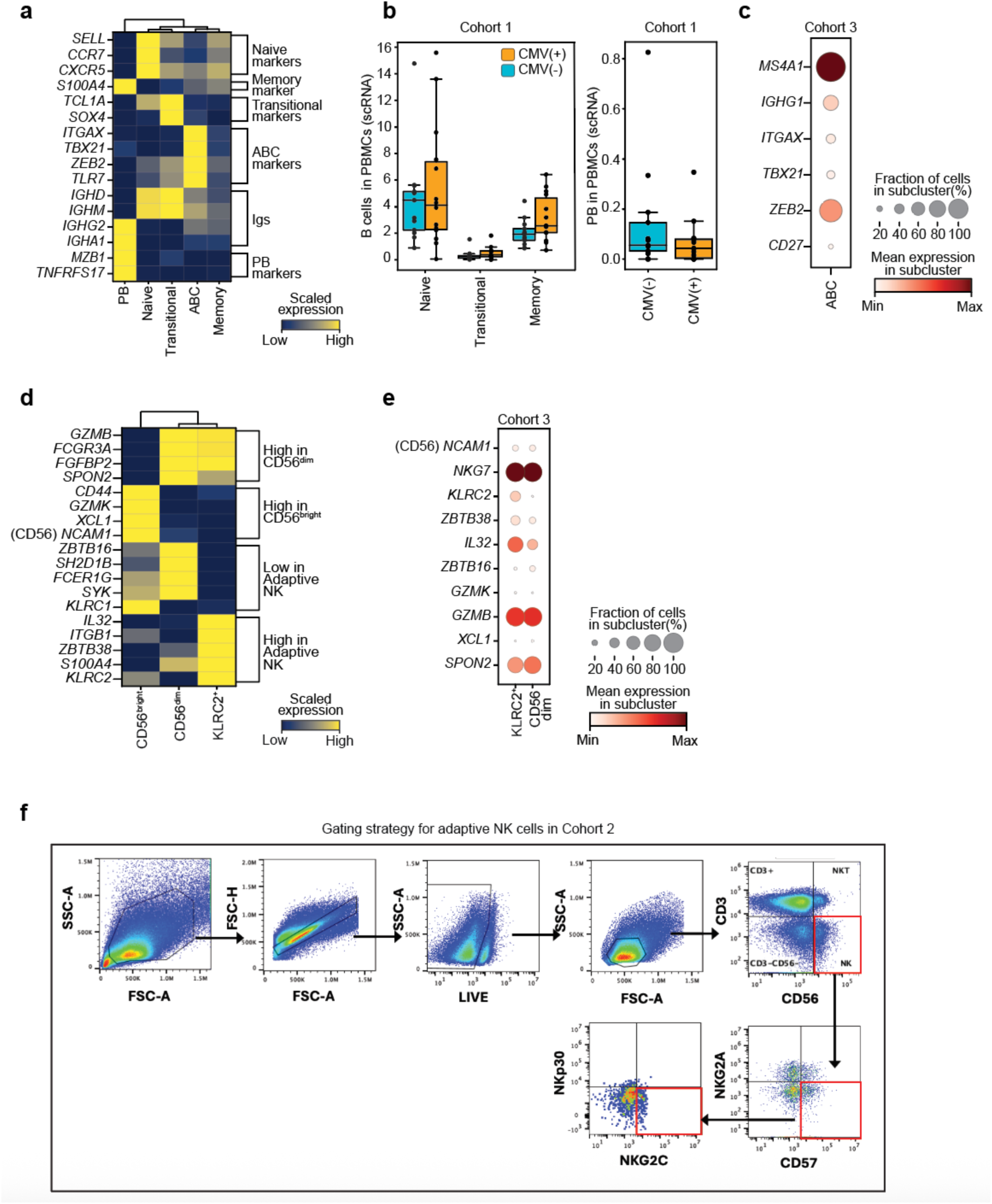
B and NK cell annotations. **a,** Scaled expression of select marker genes in B cell subsets. **b,** *Left:* naïve, transitional, and memory B cell frequencies within PBMCs. *Right:* PB cell frequencies within PBMCs. All panels represent scRNA-seq data from Discovery Cohort: CMV(-) (n=13) and CMV(+) (n=14). **c,** Scaled expression of select marker genes for ABCs in Cohort 3. **d,** Scaled expression of select marker genes in NK cell subsets. **e,** Scaled expression of select marker genes for KLRC2^+^ and CD56^dim^ NK cells in Cohort 3. **f,** Gating strategy for adaptive NK cells in Cohort 2 using flow cytometry; FSC, forward scatter; SSC, side scatter. Box plots display the median and IQR (25–75%), with whiskers representing the upper and lower quartiles ±1.5× IQR. Each dot represents an individual sample. Mann-Whitney-U test (two-sided) was used to compare cell counts between CMV(-) and CMV(+) (**b**). None of the measurements were significant after adjusting the *P*-values for multiple hypothesis testing, using Benjamini-Hochberg at false discovery rate (FDR) < 0.05.

**Extended Data Fig. 5:**
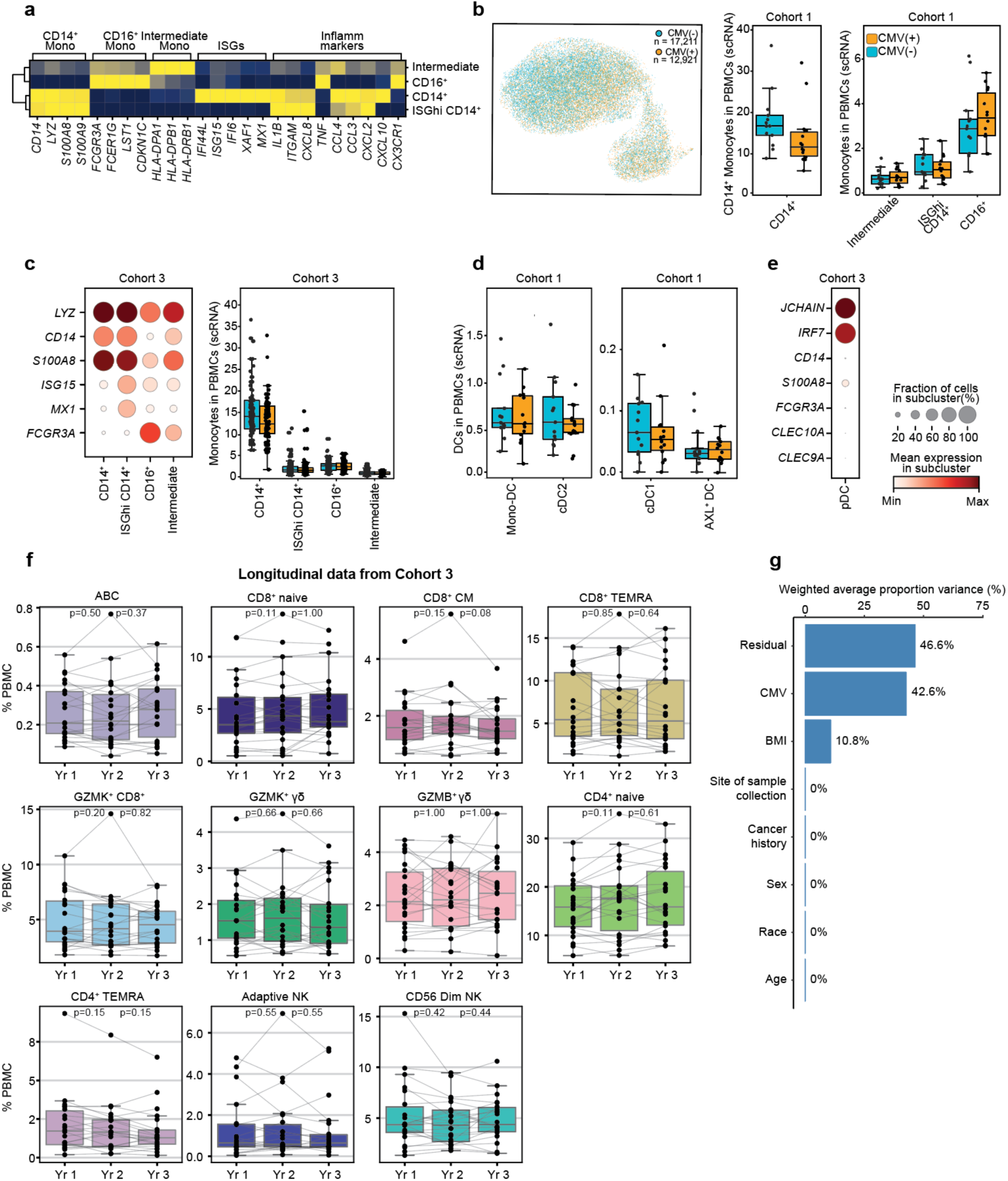
Monocyte subset annotations and temporal stability of PBMC frequencies associated with CMV. **a,** Scaled expression of select marker genes in monocyte subsets. **b,** *Left:* UMAP color coded for CMV serostatus. *Middle:* CD14^+^ monocyte frequencies within PBMCs. *Right:* Intermediate, ISGhi CD14^+^, and CD16^+^ monocyte frequencies within PBMCs. All panels represent scRNA-seq data from Discovery Cohort: CMV(-) (n=13) and CMV(+) (n=14) **c,** *Left:* scaled expression of select marker genes for CD14^+^, ISGhi CD14^+^, and CD16^+^, and intermediate monocytes. *Right:* monocyte subset frequencies within PBMCs. All panels represent scRNA-seq data from Cohort 3: CMV(+) (n=42) and CMV(-) (n=50). **d,** *Left:* Mono-DC, and cDC2 frequencies within PBMCs. *Right:* cDC1 and AXL^+^ DC frequencies within PBMCs. All panels represent scRNA-seq data from Discovery Cohort: CMV(-) (n=13) and CMV(+) (n=14). **e,** Scaled expression of select marker genes for pDCs in Cohort 3. **f,** Individual-level frequencies of CMV-associated cell types over a three-year period in CMV(+) individuals from Cohort 3 (n=22). Connecting lines represent individual-specific changes in cell frequencies between Year 1, Year 2, and Year 3 visits. *P*-values in each box plot are shown for **i,** Year 1-Year 2 and (ii) Year 2-Year 3 comparisons. **g**, Principal Variance Component Analysis (PVCA) of lymphocyte subset frequencies, estimating the proportion of variance explained by each clinical factor in Cohort 1. Box plots display the median and IQR (25–75%), with whiskers representing the upper and lower quartiles ±1.5× IQR. Each dot represents an individual sample. Mann-Whitney-U test (two-sided) was used to compare cell counts (**b**, **d, f**). None of the measurements were significant after adjusting the *P*-values for multiple hypothesis testing, using Benjamini-Hochberg at false discovery rate (FDR) < 0.05.

**Extended Data Fig. 6:**
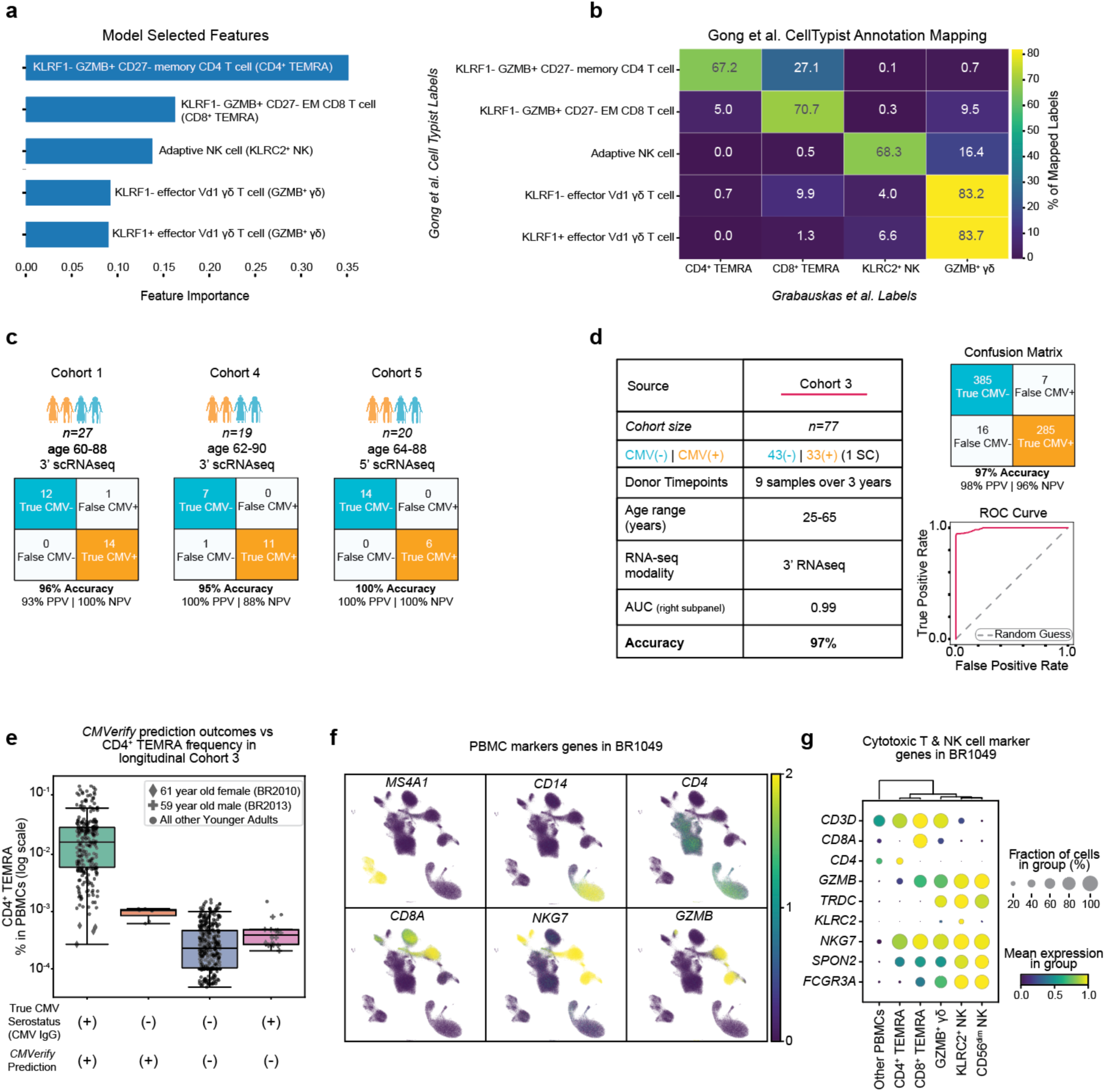
*CMVerify* selected features and extended prediction metrics. **a,** Top 5 selected features by *CMVerify* during the training procedure (see Methods) to predict CMV serostatus. Cell type labels were determined via the AIFI_L3 CellTypist model.^32^ **b,** Heatmap showing percent of cell type labels mapped from AIFI_L3 CellTypist model to manual labels in our study. **c,** Confusion matrices for CMV serostatus prediction in each independent cohort from **Fig. 5b**. **d,** *Left*: table summarizing the longitudinal Cohort 3. ‘1 SC’ stands for ‘one seroconverter’ (donor ID: BR1049). *Right top*: Confusion matrix for CMV serostatus predictions from the longitudinal Cohort 3 in **Fig. 5d**. *Right lower*: Receiver Operating Characteristic (ROC) curve quantifying performance of *CMVerify* on the longitudinal Cohort 3 in **Fig. 5d**. **e,** CMVerify prediction outcomes and CD4⁺ TEMRA frequencies per sample in the longitudinal Cohort 3. Diamonds represent the 61-year-old female (donor ID: BR2010), misclassified at five timepoints. Crosses represent the 59-year-old male (donor ID: BR2013), misclassified at nine timepoints. Circles represent all other samples. **f**, Expression of PBMC marker genes in donor BR1049 across all timepoints shown in **Fig. 5e. g,** Expression of cytotoxic marker genes in select PBMC subsets from **Fig. 5f**. PPV, positive predictive value; NPV, negative predictive value (**c,d**).

**Extended Data Fig. 7:**
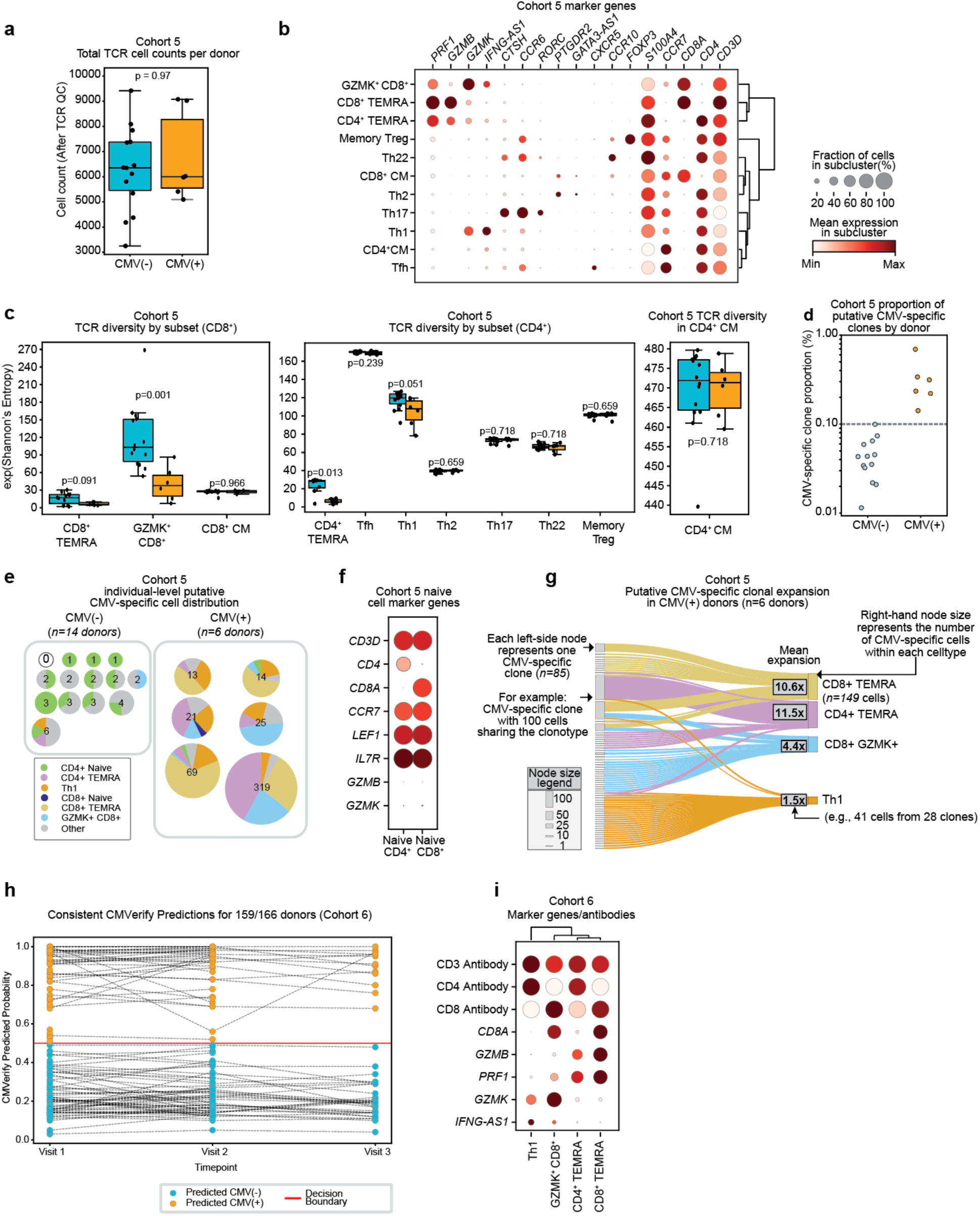
TCR diversity and CMV-specific clonal expansions in Cohorts 2 and 3. **a,** Total TCR cell counts after quality control in Cohort 2. **b,** Expression of marker genes in TCR-profiled cells in Cohort 2. **c,** TCR diversity in CD8^+^ and CD4^+^ T cells in Cohort 2. **d,** Putative CMV-specific clone proportion per donor in Cohort 2. **e,** Pie charts showing putative CMV-specific cell types per donor in CMV(–) and CMV(+) groups. Each pie chart corresponds to a donor in Cohort 2, where the sizerepresents the number of CMV-specific cells detected (log scale). One CMV(-) donor had no putative CMV-specific cell types, indicated by ‘0.’**f**, Expression of naïve marker genes in TCR-profiled cells in Cohort 2. **g,** Sankey plot of putative CMV-specific clones and their corresponding cells in CMV(+) individuals from Cohort 2. Each node on the left represents one putative CMV-specific clone, with thenode size proportional to the number of cells sharing the clonotype. Nodes on the right represent annotated cell types, with node size proportional to the number of putative CMV-specific cells detected within the celltype. **h,** Longitudinal CMVerify predictions for each individual in Cohort 3. Y-axis represents model predicted probability, each bubble is a unique sample corresponding with a donor timepoint. Samples from the same donor are connected via gray lines. Based on the Decision Boundary (y = 0.5), all bubbles below the Decision Boundary (y < 0.5) represent prediction of CMV(-), vs. bubbles above the Decision boundary (y > 0.5) represent prediction of CMV(+). Bubbles are color coded for CMVerify prediction based on the prediction probability. **i**, Expression of marker genes and antibody staining in select TCR-profiled cells in Cohort 3. For **a** and **c,** box plots display the median and IQR (25–75%), with whiskers representing the upper and lower quartiles ±1.5× IQR. Each dot represents an individual sample. Mann-Whitney-U test (two-sided) was used to compare differences between groups. For **c**, TCR diversity was calculated using Shannon’s entropy (e^H^). Additionally, due to the small sample size in the CMV(+) group (6/20), diversity calculations within cell types were not FDR-corrected.

